# A Trio of Ubiquitin Ligases Sequentially Drive Ubiquitylation and Autophagic Degradation of Dysfunctional Yeast Proteasomes

**DOI:** 10.1101/2021.05.12.443936

**Authors:** Richard S. Marshall, Richard D. Vierstra

## Abstract

As central effectors of ubiquitin (Ub)-mediated proteolysis, proteasomes are regulated at multiple levels, including degradation of unwanted or dysfunctional particles via autophagy (termed proteaphagy). In yeast, inactive proteasomes are exported from the nucleus, sequestered into cytoplasmic aggresomes via the Hsp42 chaperone, extensively ubiquitylated, and then tethered to the expanding phagophore by the autophagy receptor Cue5. Here, we demonstrate the need for ubiquitylation driven by the trio of Ub ligases (E3s) San1, Rsp5 and Hul5, which, together with their corresponding E2s, work sequentially to promote nuclear export and Cue5 recognition. Whereas San1 functions prior to nuclear export, Rsp5 and Hul5 likely decorate aggresome-localized proteasomes in concert. Ultimately, topologically complex Ub chain(s) containing both K48 and K63 Ub-Ub linkages are assembled, mainly on the regulatory particle, to generate autophagy-competent substrates. As San1, Rsp5, Hul5, Hsp42, and Cue5 also participate in general proteostasis, proteaphagy likely engages an essential mechanism for eliminating inactive/misfolded proteins.

**HIGHLIGHTS:**

- Ubiquitylation is essential for the autophagic turnover of dysfunctional proteasomes.
- The San1, Rsp5 and Hul5 E3s act sequentially to drive proteaphagy.
- The E2s Ubc1, Ubc4 and Ubc5 are collectively required.
- Both K48- and K63-mediated Ub-Ub linkages are assembled for efficient proteaphagy.

## INTRODUCTION

Autophagy and the ubiquitin (Ub)-proteasome system (UPS) are the two main protein quality control (PQC) pathways in eukaryotes that eliminate unwanted or aberrant proteins and protein complexes whose removal is essential for maintaining a healthy proteome (Marshall and Vierstra, 2019; Pohl and Dikic, 2019). In fact, situations that compromise or overwhelm their proteolytic capacities often induce proteotoxic stress, ultimately leading to the appearance of aberrant protein aggregates that become cytotoxic if allowed to accumulate. As these aggregates are emerging as hallmarks of aging, cancer, and various proteinopathies of medical relevance, including Alzheimer’s, Parkinson’s, Huntington’s and amyotrophic lateral sclerosis (Hipp et al., 2019; Rape, 2018), understanding how autophagy and the UPS converge to clear amyloidogenic proteins and maintain cellular proteostasis is therapeutically relevant.

Whereas autophagy acts by sequestering unwanted cytoplasmic material within vesicles for eventual deposition and breakdown in vacuoles (yeast and plants) or lysosomes (animals) (Marshall and Vierstra, 2018a; Reggiori and Klionsky, 2013), the UPS selectively tethers chains of Ub to appropriate substrates, which then direct their breakdown by the proteasome, an ATP-dependent, 26S protease complex (Greene et al., 2020). Autophagic specificity is guided by a suite of receptors that simultaneously recognize the substrate and the lipidated Atg8 protein that embeds within the engulfing phagophore membrane (Johansen and Lamark, 2020; Kirkin and Rogov, 2019). By contrast, the UPS engages a Ub-activating (E1), Ub-conjugating (E2) and Ub ligase (E3) reaction cascade, with a highly polymorphic set of E3s devoted to substrate recognition. There are ∼90 E3s in yeast (*Saccharomyces cerevisiae*) (Finley et al., 2012) and well over 1,500 in other species such as plants (Vierstra, 2009), thus providing widespread influence over proteomes. The ubiquitylated species are then recognized by a small set of proteasome-associated Ub receptors (Martinez-Fonts et al., 2020). In some situations, autophagy also participates in ubiquitylated protein removal using receptors with affinity for the bound Ub moieties as well as Atg8 (Pohl and Dikic, 2019; Yin et al., 2020).

At the nexus of the UPS is the proteasome, a 64 or more subunit particle assembled from two subcomplexes, the 20S core protease (CP) and the 19S regulatory particle (RP) (Greene et al., 2020). The CP compartmentalizes the proteolytic active sites within a four-ringed barrel, whereas the RP houses Ub receptors, deubiquitylating enzymes (DUBs) that recycle the Ub moieties, and a ring of AAA-ATPases that couples ATP hydrolysis to substrate unfolding and translocation into the CP lumen. Given that many substrates are nuclear, most proteasomes can be found within this compartment (Marshall et al., 2016; Pack et al., 2014). Not surprisingly, proteasome activity is controlled at multiple levels, including dedicated transcription factors that co-ordinate expression to meet demand, chaperone-mediated assembly, ATP availability that promotes CP/RP binding, post-translational modifications, and the association of various regulatory factors (Marshall and Vierstra, 2019).

Of interest here is the tight control on proteasome abundance by autophagy through an evolutionarily conserved process called proteaphagy (Cohen-Kaplan et al., 2016; Marshall et al., 2015; 2016; Waite et al., 2016). In yeast, proteaphagy occurs rapidly in response to nitrogen (N) limitation, presumably to help augment the supply of amino acids. This turnover involves nuclear export and vacuolar delivery via a bulk autophagy route activated by the nutrient-responsive Atg1 kinase (Marshall et al., 2016; Nemec et al., 2017; Waite et al., 2016). Remarkably, during carbon starvation, proteaphagy is bypassed by sequestering the CP and RP subcomplexes into cytoplasmic proteasome storage granules (PSGs) protected from breakdown (Marshall and Vierstra, 2018b). After return to carbon-replete conditions, PSGs rapidly dissolve and the RPs and CPs reassemble as nuclear 26S particles, thus restoring UPS capacity.

When impaired, proteasomes also become targets of a second proteaphagic route that removes dysfunctional species (Choi et al., 2020; Marshall et al., 2015; 2016). In yeast, compromised particles leave the nucleus and accumulate with the help of the Hsp42 chaperone into cytoplasmic aggresome puncta distinct from PSGs, which likely house other amyloidogenic proteins (Marshall et al., 2016; Peters et al., 2015). At some point, proteasomes become extensively ubiquitylated and are detected by the Cue5 autophagic receptor that simultaneously recognizes the Ub moieties through a Ub-binding CUE domain and Atg8 via an Atg8-interacting motif (AIM), thus tethering the particles to engulfing phagophores (Marshall et al., 2016). At present, the identities of the UPS component(s) that drive this ubiquitylation are unknown, and the connection between Ub and proteaphagy remains correlative. As Cue5 and its human ortholog Tollip also help clear amyloidogenic proteins of medical relevance (Lu et al., 2014), understanding the events that drive proteaphagy might help appreciate proteostasis more broadly, especially the link between protein aggregation and ubiquitylation.

Here, we investigated the connections between proteasome ubiquitylation and autophagy and demonstrated that Ub addition is essential for degrading inactivated particles, but not those removed during N starvation. From a comprehensive screen of yeast mutants compromising the UPS, we identified a trio of E3s – San1, Hul5 and Rsp5 – along with their corresponding E2s that appear to work sequentially to promote proteasome nuclear export, aggregation, and autophagy recruitment. All three E3s have been implicated in PQC (Crosas et al., 2006; Fang et al., 2011; 2014; Lu et al., 2014), suggesting that they represent core machinery for ameliorating proteotoxic stress. Further analysis of proteasome ubiquitylation detected both K48- and K63-linked Ub-Ub polymers, suggesting that a complex Ub chain architecture is needed to generate autophagy-competent substrates suitable for Cue5 recognition.

## RESULTS

### Ubiquitylation is Essential for Proteaphagy of Inactive Proteasomes

As a first step in confirming the importance of ubiquitylation to inhibitor-induced proteaphagy, we exploited a temperature-sensitive (*ts*) allele (*uba1-204*) impacting the single yeast E1 (Ghaboosi and Deshaies, 2007). While this Uba1 variant is active in cells grown at the permissive temperature of 30°C, it is rapidly compromised upon switching cells to the non-permissive temperature of 37°C, which induces a precipitous loss of Ub conjugates. By applying the “free GFP release assay” in combination with yeast strains in which the CP and RP subunits Pre10 (α_7_) and Rpn5, respectively, are tagged with GFP (Marshall et al., 2016), we assayed for delivery of proteasomes to vacuoles upon N starvation or inactivation by the proteasome inhibitor MG132. As demonstrated for numerous substrates (*e.g*., Lu et al., 2014; Marshall et al., 2016; 2019; Lee et al., 2020), autophagy of such fusions causes rapid breakdown of the tagged protein concomitant with the accumulation of the more stable free GFP moiety inside vacuoles, the appearance of which can be easily detected by immunoblotting with anti-GFP antibodies.

When first testing the impact of N starvation by the free GFP release assay, we grew log-phase WT and *uba1-204* cells expressing Pre10-GFP or Rpn5-GFP at 30°C in +N medium, and then either kept them in +N medium, switched them to –N medium at 30°C for 8 hr, or exposed them to 37°C for 1 hr to inactivate Uba1-204 followed by maintenance on +N medium or a switch to –N medium for 8 hr at 37°C. As shown in Figures 1A and 1B, proteasomes were stable in WT when grown in +N medium at 30°C or 37°C, but were rapidly degraded upon transfer to –N medium (as measured by the appearance of free GFP) via a mechanism absent in autophagy-deficient *Δatg7* cells (Marshall et al., 2016; Waite et al., 2016). When the *uba1-204* cells were similarly tested in –N medium, the appearance of GFP was evident both at 30°C and 37°C, demonstrating that ubiquitylation is not required for proteaphagy during N starvation.

**Figure 1.**
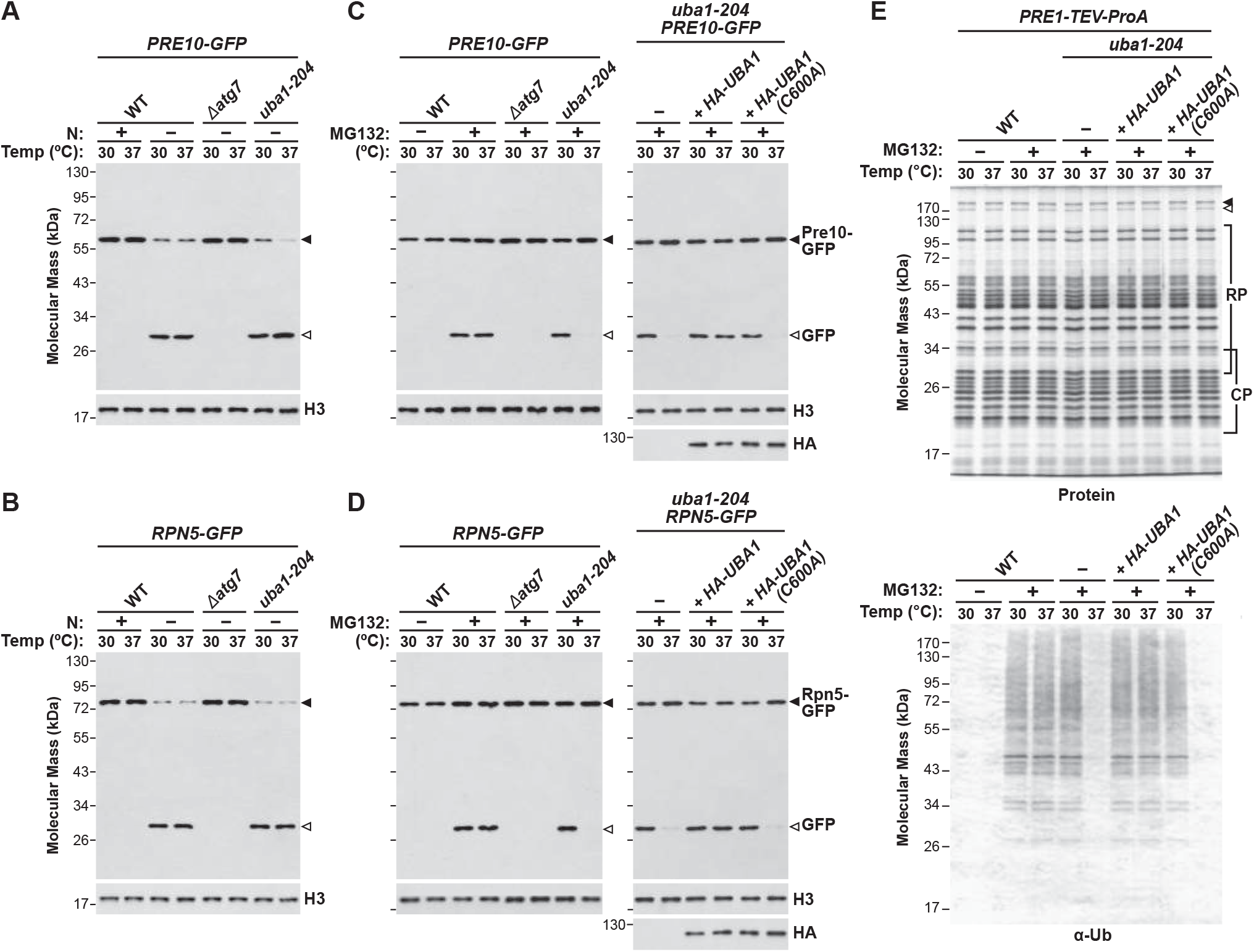
Ubiquitylation is Essential for the Autophagic Degradation of Inhibited Yeast Proteasomes. **(A and B)** Proteaphagy proceeds normally in *uba1-204* cells during N starvation at the permissive (30°C) and non-permissive (37°C) temperatures, as measured by the free GFP release assay. WT and *uba1-204* cells expressing *PRE10-GFP* **(A)** or *RPN5-GFP* **(B)** grown on +N medium at 30°C were either kept on +N medium or switched to –N medium and grown for 8 hr at either 30°C or 37°C. Release of free GFP was assayed by immunoblotting cell extracts with anti-GFP antibodies. Anti-histone H3 antibodies confirmed near equal protein loading. The *Δatg7* mutant was included as a positive control. **(C** and **D)** Proteaphagy induced by MG132 is blocked at the non-permissive temperature in *uba1-204* cells. *PRE10-GFP* **(C)** and *RPN5-GFP* **(D)** cells, either WT or harboring the *uba1-204* mutation with or without HA-tagged wild-type Uba1 or the Uba1(C600A) active site mutant, were treated with or without 80 µM MG132 for 8 hr at either 30°C or 37°C. Release of free GFP was assayed as in (A). Anti-HA antibodies confirmed expression of HA-tagged Uba1. **(E)** Inactivation of Uba1 blocks ubiquitylation of inhibited proteasomes. *PRE1-TEV-ProA* cells, either WT or harboring the *uba1-204* mutation with or without HA-tagged versions of Uba1 or Uba1(C600A), were grown as in (C). Affinity-purified proteasomes were subjected to SDS-PAGE and either stained for protein or immunoblotted for Ub. CP and RP subunits are highlighted by the brackets. Closed and open arrowheads locate the Blm10 and Ecm29 accessory proteins, respectively.

To examine MG132-inhibited proteasomes, inhibitor uptake was enabled by performing all experiments in the *Δerg6* background that increases drug permeability (Lee and Goldberg, 1996). While proteasomes were rapidly degraded in WT *PRE10-GFP* or *RPN5-GFP* cells exposed to MG132 either at 30°C and 37°C, as judged by the appearance of free GFP, they were stable in the *Δatg7* background at either temperature (Figures 1C and 1D; Marshall et al., 2016). By contrast, when the *uba1-204* background was tested, inhibited proteasomes were rapidly degraded at 30°C but were stable at 37°C. For further proof that the E1 activity of Uba1 was required for this turnover, we complemented the *uba1-204* allele with HA-tagged versions of either wild-type Uba1 or the Uba1(C600A) active site mutant (Lee and Schindelin, 2008). As shown in Figures 1C and 1D, full rescue of free GFP release was seen when HA-Uba1 was introduced into *uba1-204* cells treated with MG132 at 37°C, but not when HA-Uba1(C600A) was introduced instead.

To confirm a block in proteasome ubiquitylation, we monitored the ubiquitylation status of 26S particles affinity purified from *uba1-204* cells grown with or without MG132 at 30°C or 37°C, including those complemented with HA-Uba1 or HA-Uba1(C600A). Here, a purification strain developed by Leggett *et al*. (2002) was employed in which the Pre1 (β_4_) subunit was replaced by a TEV-cleavable Protein A (ProA)-tagged version suitable for affinity enrichment with IgG beads. The resulting preparations displayed the characteristic ladder of CP and RP subunits upon SDS-PAGE when isolated in the presence of ATP, and included the Blm10 and Ecm29 accessory proteins that migrate above 130 kDa (Figure 1E). Importantly, the *uba1-204* allele, MG132 exposure, and/or growth at 30°C and 37°C or did not appreciably alter the subunit profile of the 26S particles. However, when immunoblotted for Ub, strong proteasome ubiquitylation was evident in WT upon treatment with MG132 either at 30°C or 37°C (Figure 1E). High salt washes failed to remove this signal, implying that it reflected ubiquitylated proteasomes as opposed to ubiquitylated substrates remaining attached during purification (Marshall et al., 2016; Peth et al., 2010). When proteasomes were similarly purified and analyzed from *uba1-204* cells, strong ubiquitylation was evident when the MG132-treated cells when grown at 30°C, but not when grown at 37°C (Figure 1E). As expected, introduction of HA-Uba1 in the *uba1-204* background restored Ub addition at 37°C, while the HA-Uba1(C600A) mutant failed, thus confirming that an active E1 is required for this modification.

As a final step in demonstrating the need for ubiquitylation during inhibitor-induced proteaphagy, we monitored the autophagic transport of GFP-tagged proteasomes by fluorescence confocal microscopy. As seen previously (Marshall et al., 2016; Pack et al., 2014), most GFP fluorescence in WT *PRE10-GFP* cells grown with N at either 30°C or 37°C resided in the nucleus (Figures 2A and 2C). When N starved, the GFP signal from WT and *uba1-204* cells then concentrated in vacuoles at both 30°C or 37°C (Figure S1). However, when WT cells were exposed to MG132, while most of this signal quantified morphometrically again left the nucleus, it initially became concentrated into cytoplasmic puncta, assigned here as aggresomes by their co-localization with the aggresome marker Rnq1-mCherry (Marshall et al., 2016; Peters et al., 2015), before ending up in vacuoles (Figures 2A and 2C). In fact, by 8 hr most GFP signal was vacuolar, consistent with the autophagic clearance of inhibited proteasomes as seen by the free GFP release assay. Strikingly, this transport was still evident in MG132-treated *uba1-204* cells maintained at 30°C, but was substantially lost at 37°C, with proteasomes instead remaining concentrated in nuclei. As above, this vacuolar deposition required active E1. Whereas the *uba1-204* mutant complemented with HA-Uba1 restored accumulation within aggresomes and subsequent vacuolar deposition when grown at 37°C, this movement was not restored by the HA-Uba1(C600A) mutant (Figures 2B and 2C). Taken together, our studies on the *uba1-204* mutant not only revealed that proteaphagy of inhibited proteasomes requires prior ubiquitylation, but also demonstrated that their nuclear export depends on this modification.

**Figure 2.**
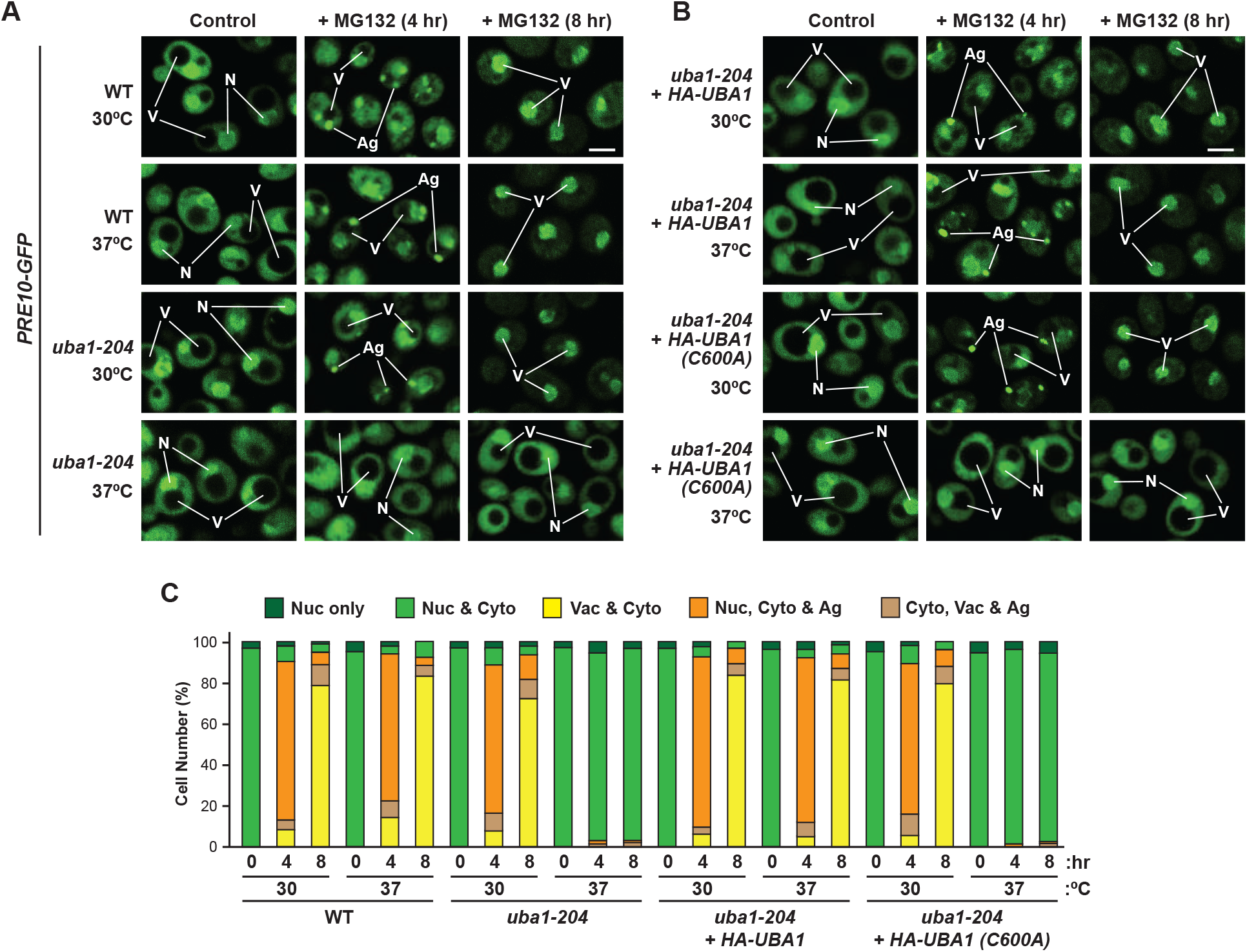
Ubiquitylation Helps Deliver Inhibited Proteasomes to Vacuoles by Autophagy. **(A and B)** Inhibited proteasomes labelled with Pre10-GFP are transported to vacuoles in WT but not *uba1-204* cells at the non-permissive temperature. *PRE10-GFP* cells were grown as in Figure 1A and imaged by confocal fluorescence microscopy at 0, 4 or 8 hr after treatment with 80 µM MG132. **(A)** Comparisons of *PRE10-GFP* cells with or without the *uba1-204* mutation. **(B)** Comparisons of *PRE10-GFP uba1-204* cells complemented with HA-tagged Uba1 or the Uba1(C600A) active site mutant. N, nucleus; V, vacuole; Ag, cytoplasmic aggregate. Scale bar, 2 µm. **(C)** Quantification of the cellular distribution of proteasomes in (A) and (B). Each bar represents the analysis of at least 200 cells.

### The E3s Hul5, Rsp5 and San1 Direct Proteasome Ubiquitylation during Proteaphagy

Having confirmed that ubiquitylation is essential for the proteaphagy of inactive particles, we next set out to genetically identify the relevant E3(s). Here, we screened by the free GFP release assay an extensive collection of yeast UPS mutants within the Targeted Ubiquitin System (TUS) deletion library developed by Hochstrasser and colleagues (Hickey et al., 2021; Ravid and Hochstrasser, 2007), which we supplemented with available *ts* alleles for essential loci. For the E3s, this collection covered 81 of the ∼90 known E3 types, and included essential subunits for multi-subunit E3s such as the anaphase-promoting complex (Apc11 and Cdc20) and the cullin-RING ligases (Cdc53, Rtt101, Cul3, and Skp1) (Finley et al., 2012). To perform the screen, each strain was modified to include the *Δerg6* mutation before mating with strains expressing the *PRE10-GFP* or *RPN5-GFP* reporters. Following sporulation, the correct strains were confirmed by PCR-based genotyping and fluorescence microscopic detection of the GFP tag. We grew each strain for 8 hr at 30°C or 37°C (as appropriate) with MG132, and then assayed for the appearance of free GFP by immunoblot analysis of culture lysates. This simple assay proved to be remarkably robust, and could measure even modest reductions in proteaphagy after MG132 treatment (Figures S2A and S2B).

From analysis of the E3 deletion strains with the Pre10-GFP reporter, we identified the HECT E3 Hul5 and the RING E3 San1 as important for inhibitor-induced proteaphagy, which we then confirmed by comparable analyses with the Rpn5-GFP reporter (Figure 3A; Figure S2A). While both deletion strains retained modest proteasome turnover in the presence of MG132 (as judged by free GFP levels), this turnover was absent in the *Δhul5 Δsan1* double mutant, suggesting that the two E3s are non-redundant. Neither the *Δhul5* and *Δsan1* single mutants or the double mutant suppressed proteaphagy induced by N starvation (Figure S3A), nor did they impair bulk autophagy as seen by Pho8Δ60 assays that measure autophagic flux by vacuolar activation of the Pho8 phosphatase (Noda and Klionsky, 2008) (Figure S4A), demonstrating that bulk autophagy was not compromised. To confirm that the ligase activities of Hul5 and San1 were essential, we complemented the corresponding deletion strains with HA-tagged wild-type or mutant versions impacting residues key to catalysis. For Hul5, C878, which forms the E3-Ub adduct prior to transfer (Crosas et al., 2006), was replaced with an alanine, while for San1, C257, which constitutes part of the RING domain critical for binding the E2-Ub adduct (Gardner et al., 2005), was replaced with a serine. As shown in Figures S3B and S3C, both wild-type HA-Hul5 and HA-San1 effectively rescued proteaphagy in the corresponding *Δhul5* and *Δsan1* strains, but this rescue failed when the HA-Hul5(C878A) and HA-San1(C257S) mutants were used instead.

**Figure 3.**
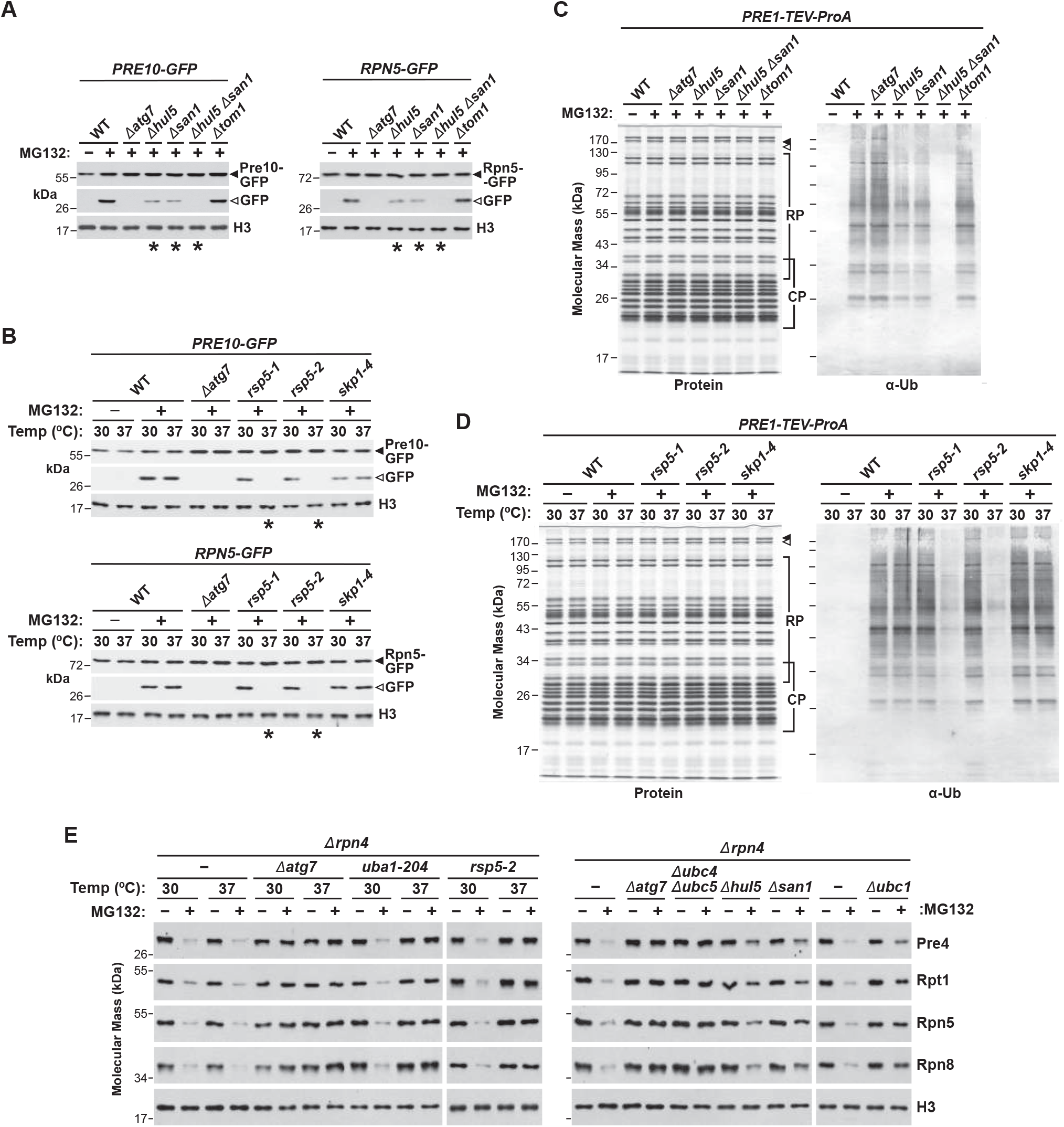
A Trio of Ub Ligases (E3s) Promote the Ubiquitylation and Autophagic Degradation of Inhibited Proteasomes. **(A)** Hul5 and San1 deletion mutants dampen proteaphagy of inhibited proteasomes, as assayed by the free GFP release assay. WT and mutant cells expressing Pre10-GFP (left panels) or Rpn5-GFP (right panels) were grown for 8 hr at 30°C on +N medium with or without 80 µM MG132 and assayed for free GFP as in Figure 1A. Only the sections of the gels harboring the GFP fusion and free GFP are shown. The unrelated *Δtom1* E3 mutant was included as a negative control. See Figures S2A and 2B for the full screen of yeast E3 mutants. **(B)** *ts* alleles impacting Rsp5 dampen proteaphagy of inhibited proteasomes. WT and mutant cells expressing *PRE10-GFP* (top panels) or *RPN5-GFP* (bottom panels) were grown for 8 hr at 30°C or 37°C on +N medium with or without MG132, and assayed as in (A). The *ts* allele of the unrelated E3 component Skp1 was included as a negative control. **(C** and **D)** Hul5 and San1 **(C)** or Rsp5 **(D)** mutants dampen ubiquitylation of inhibited proteasomes. *PRE1-TEV-ProA* cells either WT or harboring the *Δhul5*, *Δsan1*, *Δhul5 Δsan1, rsp5-1* or *rps5-2* mutations were grown with or without MG132 as in (A) or (B). Affinity-purified proteasomes were subjected to SDS-PAGE, and either stained for protein or immunoblotted for Ub, as in Figure 1E. Proteasomes from the *Δatg7*, *Δtom1* or *skp1-4* backgrounds were included as controls. **(E)** The Uba1 E1, the Ubc1, Ubc4 and Ubc5 E2s, and the Hul5, San1 and Rsp5 E3s together assist in proteaphagy of inhibited, non-tagged proteasomes. WT and mutant cells also harboring the *Δrpn4* deletion were grown with or without MG132 at 30°C or 37°C as in (A) or (B), and assayed by immunoblotting for CP (Pre4) and RP (Rpt1, Rpn5 and Rpn8) subunits. Left panel: analysis of the *uba1-204* and *rsp5-2 ts* mutants. Right panel: analysis of the *Δubc1*, *Δubc4 Δubc5*, *Δhul5* and *Δsan1* deletion mutants.

From analysis of the *ts* E3 collection by the free GFP release assay, we also identified the HECT E3 Rsp5 as essential for inhibitor-induced proteaphagy. While this turnover occurred normally in the *rsp5-1* and *rsp5-2 ts* strains (Lu et al., 2014) at 30°C, it was abolished at the non-permissive temperature of 37°C (Figure 3B; Figure S2B). Neither *rsp5* allele affected proteaphagy during N starvation at either 30°C or 37°C (Figures S3D), nor bulk autophagy as measured by the Pho8Δ60 reporter (Figure S4B), suggesting that the *ts* alleles did not compromise general autophagy. We additionally confirmed that the Ub transferase activity of Rsp5 was essential by complementing the *rsp5-2* mutant with HA-tagged wild-type or inactive F618A and C777A versions that compromised Ub binding and formation of the Ub-E3 adduct, respectively (French et al., 2009; Wang et al., 1999). Whereas wild-type HA-Rsp5 effectively rescued proteaphagy in *rsp5-2* cells when grown with MG132 at 37°C, the HA-Rsp5(F618A) and HA-Rsp5(C777A) variants failed (Figure S3E). Remarkably, Hul5, San1 and Rps5 have been previously linked to dysfunctional protein clearance more generally (Crosas et al., 2006; Fang et al., 2011; 2014), including a direct connection between Rsp5 and the autophagy receptor Cue5 (Lu et al., 2014), indicating that they might also ameliorate proteotoxic stress more broadly.

As further confirmation that Hul5, San1 and Rps5 promote the turnover of wild-type proteasomes and not just those decorated with Pre10-GFP and Rpn5-GFP, we examined total proteasome levels in the mutant backgrounds with subunit-specific antibodies (Figure 3E). Here, WT cells were modified to harbor the *Δrpn4* mutation to eliminate transcriptional up-regulation of proteasomes by proteasome inhibitors (Xie and Varshavsky, 2001), grown at 30°C or 37°C for 8 hr with MG132, and then immunoblotted with antibodies to both CP (Pre4 (β_7_)) and RP (Rpt1, Rpn5, and Rpn8) subunits. Whereas WT cells lost both the CP and RP after treatment with MG132, this loss was effectively stalled in *uba1-204* and *rsp5-2* cells grown at 37°C, as in *Δatg7* cells, and was substantially reduced in *Δhul5* and *Δsan1* cells, in line with their more modest effects on proteaphagy (Figure 3E).

Predicting that San1, Hul5, and Rsp5 ubiquitylate inhibited proteasomes, we affinity purified 26S particles via the Pre1-TEV-ProA tag from MG132-treated WT and mutant cells and assessed their ubiquitylation by immunoblotting as above. As shown in Figures 3C and 3D, the *Δhul5*, *Δsan1*, and *rsp5-2* mutations had no discernable effect on the protein profile of CP and RP subunits, nor binding of Blm10 and Ecm29 at either 30°C or 37°C. However, when samples from MG132-treated cells were immunoblotted for Ub, the signal was substantially reduced in the *Δhul5* and *Δsan1* single mutants, and nearly absent in the *Δhul5 Δsan1* double mutant and in the *rps5-2* mutant grown at 37°C (Figures 3C and 3D). Consequently, we consider it likely that Hul5, San1 and Rsp5 are the main yeast ligases driving proteasome ubiquitylation upon inhibition, even though our initial screen did not test all possibilities.

We next monitored how the three E3s might influence autophagic transport of inhibited proteasomes by fluorescence confocal microscopy and morphometric quantification of Pre10-GFP tagged particles (Marshall et al., 2016). Surprisingly, substantial differences were seen for the *Δsan1* versus *Δhul5* and *rsp5-2* backgrounds (Figures 4A-4D). Before MG132 treatment, proteasomes displayed the typical nuclear and cytoplasmic distributions in *Δhul5, Δsan1*, *Δhul5 Δsan1,* and *rsp5-2* cells at both 30°C and 37°C, indicating that the trio did not impact normal proteasome compartmentalization. However, while proteasomes in WT first coalesced into aggresomes 4 hr after MG132 exposure, and then were translocated to vacuoles by 8 hr (Marshall et al., 2016), they remained mostly nuclear in *Δsan1* cells, with little to no GFP signal appearing in cytoplasmic puncta or vacuoles (Figures 4A and 4C). By contrast, inhibited proteasomes in *Δhul5* and *rsp5-2* cells exited the nucleus and coalesced into aggresomes after 4 hr, but failed to accumulate in vacuoles even after 8 hr. Instead, these puncta appeared to grow larger, which for *rps5-2* cells eventually became the dominant GFP signal after longer-term proteasome inhibition at 37°C (Figure 4C). Notably, proteasomes remained nuclear in the *Δhul5 Δsan1* double mutant, implying that San1 is epistatic to Hul5 (Figures 4A and 4C).

**Figure 4.**
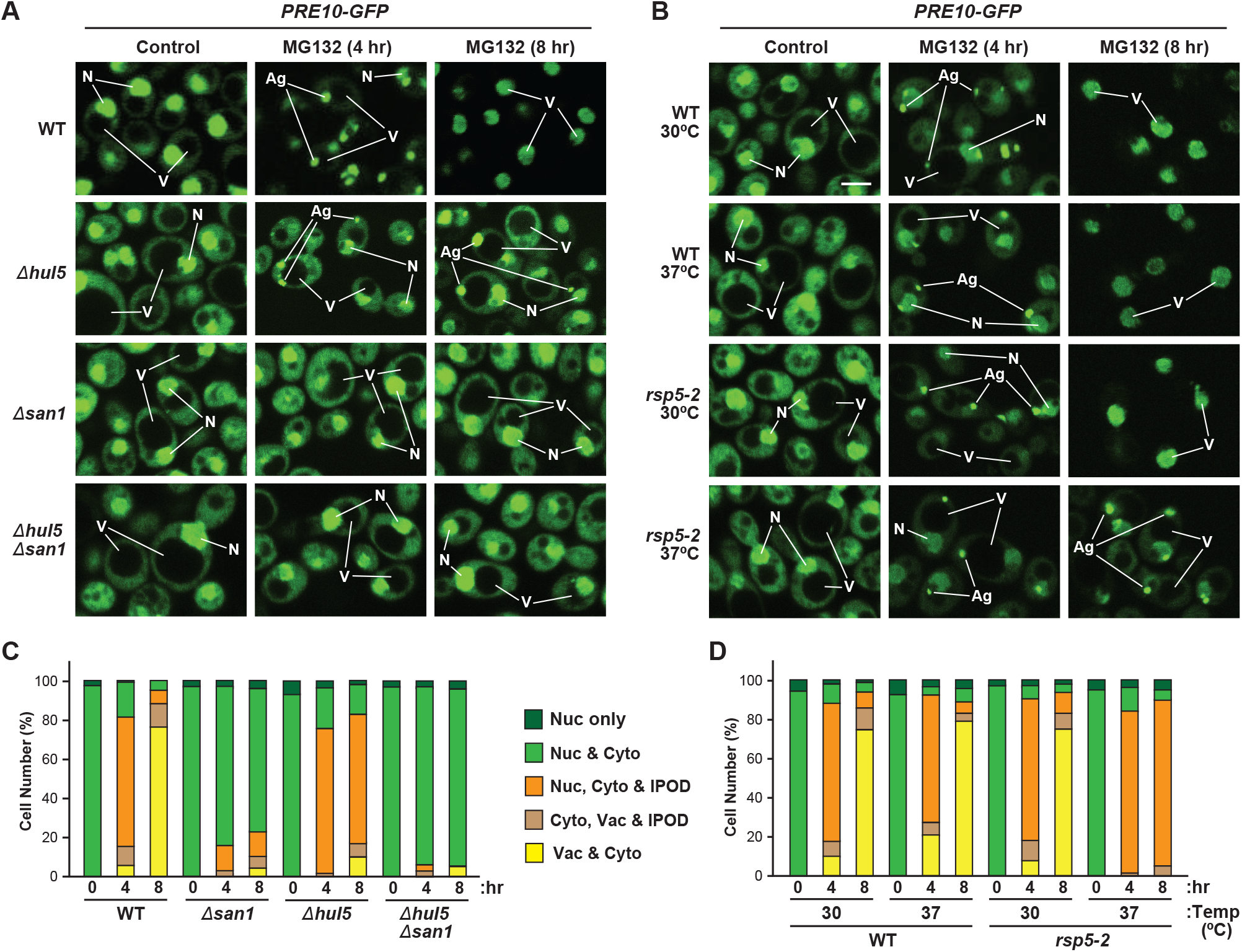
The Hul5, San1, and Rsp5 E3s Have Distinct Impacts on Proteaphagy of Inhibited Proteasomes. **(A)** Inhibited proteasomes are not transported to vacuoles and either condense into cytoplasmic aggregates in *Δhul5* cells, or are retained in the nucleus in *Δsan1* and *Δhul5 Δsan1* cells. WT and mutant *PRE10-GFP* cells were grown at 30°C and imaged by confocal fluorescence microscopy 0, 4 and 8 hr after MG132 treatment, as in Figure 2A. **(B)** Inhibited proteasomes are not transported to the vacuole in *rsp5-2* cells at the non-permissive temperature and instead condense into cytoplasmic aggresomes. WT and mutant *PRE10-GFP* cells were grown at 30°C or 37°C with or without MG132 and then imaged by confocal fluorescence microscopy as in (A). **(C** and **D)** Quantification of the cellular distribution of proteasomes in (A) and (B). Each bar represents the analysis of at least 200 cells.

As expected, all defects in the vacuolar delivery of proteasomes in *Δhul5*, *Δsan1*, and *rsp5-2* cells were rescued by introducing wild-type versions of HA-Hul5, HA-San1 or HA-Rsp5, respectively, but not by introducing the HA-Hul5(C878A), HA-San1(C257S), HA-Rsp5(F618A) or HA-Rsp5(C777A) variants (Figures S5A-5D). Taken together, we propose that San1 operates upstream of Hul5 and Rsp5 and speculate that it drives a ubiquitylation event that helps transport inhibited proteasomes out of the nucleus, while Hul5 and Rsp5 are needed for cytoplasmic ubiquitylation events that help transition inhibited proteasomes from aggresomes into vacuoles in concert with Cue5.

### The E2s Ubc1, Ubc4 and Ubc5 Are Also Required for Proteaphagy

The myriad of E3s work selectively with dedicated E2 isoforms that donate activated Ub as a thioester-linked E2-Ub adduct. To identify which of the 12 E2s (Ubc1-Ubc8 and Ubc10-Ubc13) or two known E2 variants (Mms2 and Stp22/Vps23) (Finley et al., 2012) are required for inhibited proteasome ubiquitylation and turnover along with Hul5, San1, and Rsp5, we screened the yeast E2 deletion collection within the TUS library, along with a viable *Δubc1* mutant in the W303 background (Gardner et al., 2005) and the *ts cdc34-2* allele for the essential E2 Cdc34/Ubc3 (Liu et al., 1995). Based on free GFP release assays using both Pre10-GFP and Rpn5-GFP at either 30°C or 37°C, we found that only Ubc1 and the functionally redundant Ubc4 and Ubc5 pair were critical (Figures 5A and 5B; Figure S2C). Strikingly, these three E2s were previously shown to work specifically with the three E3s we identified here. Ubc1 and Ubc4/Ubc5 assist San1 (Gardner et al., 2005; Ibarra et al., 2016; Matsuo et al., 2011), whereas Ubc4/Ubc5 have been connected to Hul5 and Rsp5 (Fang et al., 2011; Stoll et al., 2011).

**Figure 5.**
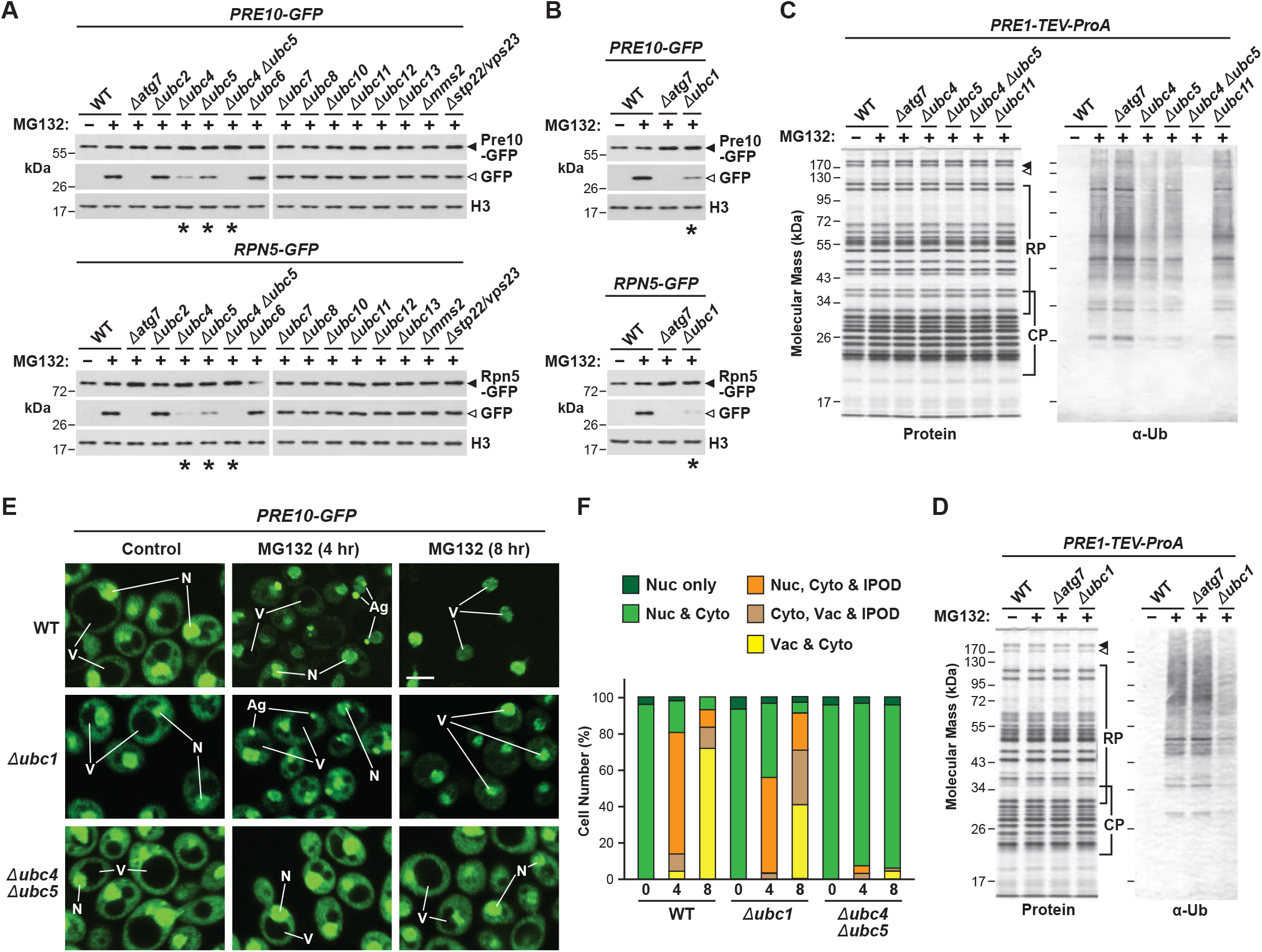
Autophagic Degradation of Inhibited Proteasomes Involves the Ubc1, Ubc4 and Ubc5 E2s. **(A and B)** Screen of a deletion mutant collection impacting 11 of the 12 known yeast E2s and two E2-like proteins by the free GFP release assay identifies Ubc4 and Ubc5 **(A)** and Ubc1 **(B)** as important for proteaphagy of inhibited proteasomes. WT and mutant cells expressing *PRE10-GFP* (top panels) or *RPN5-GFP* (bottom panels) were grown for 8 hr at 30°C on +N medium with or without 80 µM MG132, and then assayed for free GFP, as in Figure 3A. See Figure S2C for parallel analysis of a *ts* allele of the Cdc34/Ubc3 E2. **(C** and **D)** Deletion of Ubc4 and Ubc5 **(C)** or Ubc1 **(D)** dampen ubiquitylation of inhibited proteasomes. *PRE1-TEV-ProA* cells either WT or harboring the *Δubc1*, *Δubc4*, *Δubc5*, or *Δubc4 Δubc5* mutations were grown for 8 hr at 30°C with or without MG132 as in (A). Affinity-purified proteasomes were subjected to SDS-PAGE and either stained for protein or immunoblotted for Ub, as in Figure 1E. Proteasomes from *Δubc11* cells were included as a negative control. **(E)** The *Δubc1* and *Δubc4 Δubc5* mutations delay or block transport of inhibited proteasomes to the vacuole, respectively. *PRE10-GFP* cells were grown as in (A) and then imaged by confocal fluorescence microscopy 0, 4 or 8 hr after MG132 treatment, as in Figure 2A. **(F)** Quantification of the cellular distribution of proteasomes in (E). Each bar represents the analysis of at least 200 cells.

While modest rates of MG132-induced proteaphagy were seen by the accumulation of free GFP with the *Δubc1*, *Δubc4* and *Δubc5* single mutants, it was effectively abolished in the *Δubc4 Δubc5* double mutant. None of the three E2 mutants suppressed proteaphagy during N starvation (Figures S6A and S6E), nor bulk autophagy as measured by the Pho8Δ60 reporter (Figure S4A), suggesting that the effect was specific to proteaphagy of inhibited 26S particles. Again, we confirmed that the conjugating activities of Ubc1, Ubc4, and Ubc5 were essential by complementing the corresponding deletion strains with HA-tagged variants impacting the active site cysteine. Whereas the wild-type versions of each E2 rescued the corresponding deletion during inhibited proteasome turnover, the Ubc1(C88A), Ubc4(C86A) and Ubc5(C86A) cysteine-to-alanine substitutions failed (Figures S6B and S6F). We also confirmed that Ubc1 and Ubc4/Ubc5 promoted turnover of wild-type proteasomes in the presence of MG132, and not just those labeled with Pre10-GFP or Rpn5-GFP. As seen by immunoblotting *Δrpn4* cell lysates with anti-CP and RP subunit antibodies, all subunits were more abundant in the *Δubc1* and *Δubc4 Δubc5* strains versus WT upon MG132 treatment (Figure 3E).

As predicted, these E2 mutants also compromised proteasome ubiquitylation after inactivation. When compared to MG132-treated WT and *Δatg7* cells, this conjugation was substantially reduced in *Δubc1*, *Δubc4* and *Δubc5* cells, and was effectively eliminated in *Δubc4 Δubc5* cells, without an apparent impact on 26S particle composition (Figures 5C and 5D). The E2 mutants also compromised autophagic transport of proteasomes as measured by fluorescence confocal microscopy and morphometric quantification of MG132-treated *PRE10-GFP* cells (Figures 5E and 5F). The *Δubc1* cells displayed a modest suppression of proteasome transport, although the initial appearance of GFP-labelled aggresomes and final deposition of GFP fluorescence into vacuoles were still evident. By contrast, the entire route was stalled in *Δubc4 Δubc5* cells; proteasomes remained concentrated in nuclei along with a diffuse cytoplasmic fluorescence, with little to no GFP signal found in cytoplasmic puncta or in vacuoles, even after 8 hr of MG132 exposure.

### Robust Proteasome Ubiquitylation Requires Hsp42 and Preferentially Impacts the RP

Our prior studies connected Hsp42 to inhibitor-induced proteaphagy, presumably by helping coalesce proteasomes into aggresomes after exiting the nucleus (Marshall et al., 2016), which was consistent with its role in abnormal protein aggregation and clearance (Specht et al., 2011). To assess how Hsp42-mediated aggregation also influences proteasome ubiquitylation, we affinity purified Pre1-TEV-ProA-tagged 26S particles from WT and *Δhsp42* cells pre-treated with or without MG132 and measured their ubiquitylation status by immunoblotting for Ub. As shown in Figure 6A, the *Δhsp42* background did not influence the composition of proteasomes as seen by protein staining, but blocked robust ubiquitylation. This impact placed Hsp42 upstream of proteasome aggregation and the pronounced ubiquitylation steps involving Rps5 and Hul5.

**Figure 6.**
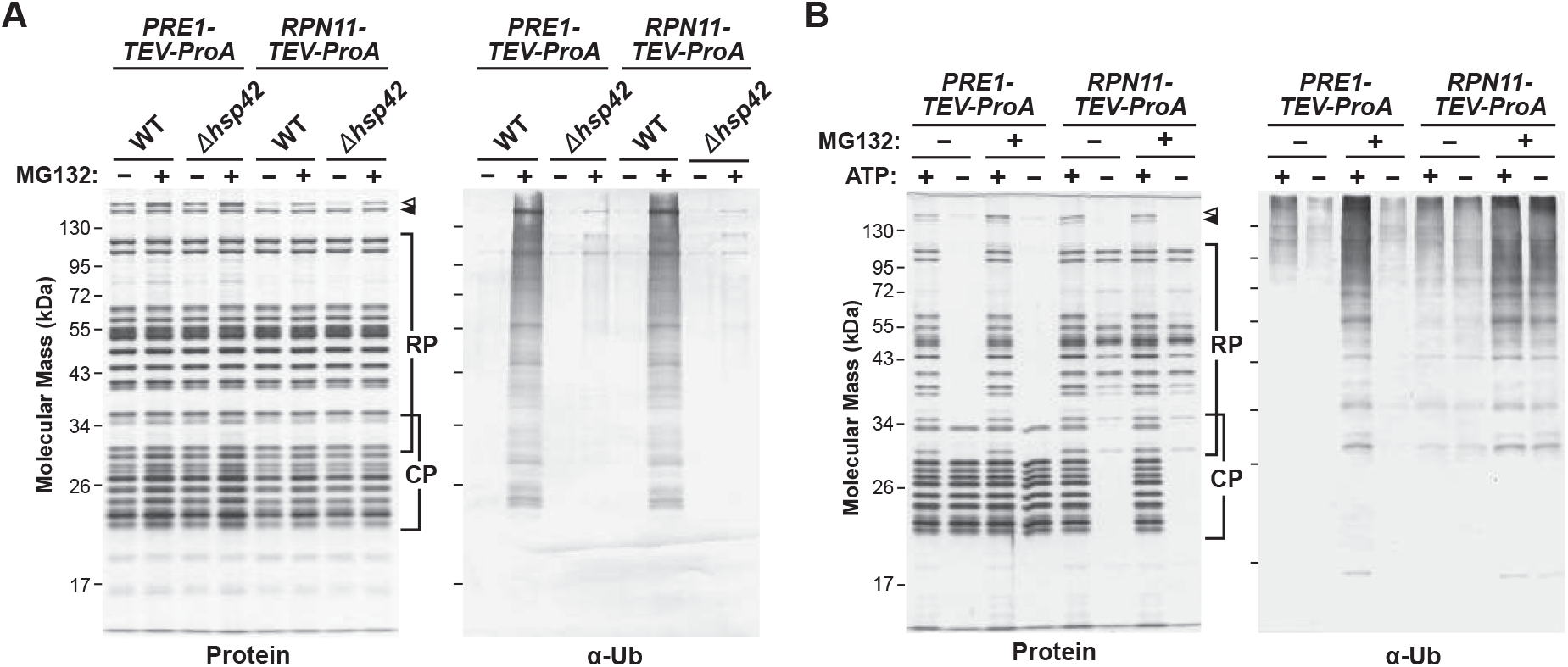
Ubiquitylation of Inhibited Proteasomes Requires Hsp42 and Preferentially Modifies the RP. **(A)** Ubiquitylation of inhibited proteasomes requires Hsp42. *PRE1-TEV-ProA* cells either WT or harboring the *Δhsp42* mutation were grown for 8 hr at 30°C with or without 80 µM MG132. Affinity-purified proteasomes were subjected to SDS-PAGE and either stained for protein or immunoblotted for Ub, as in Figure 1E. **(B)** Proteasome ubiquitylation upon inhibition mainly impacts the RP. Proteasomes were affinity purified from *PRE1-TEV-ProA* (CP) and *RPN11-TEV-ProA* (RP) cells pre-treated with or without 80 µM MG132 for 8 hr, and affinity purified either in the presence of ATP and 50 mM NaCl to promote CP/RP association, or without ATP and with 250 mM NaCl to promote CP/RP dissociation. The samples were subjected to SDS-PAGE and either stained for protein or immunoblotted for Ub as in (A).

To help locate where Ub becomes attached to proteasomes upon MG132 treatment, we exploited the fact that ATP helps maintain a strong CP/RP connection to ask whether these two subparticles were differentially modified. Here, proteasomes were affinity purified in the presence or absence of ATP from cells expressing Pre1-TEV-ProA or Rpn11-TEV-ProA that would differentially tag the CP or RP, respectively. As seen by the profile of proteasome subunits detected by protein staining, the CP was preferentially enriched from *PRE1-TEV-ProA* cells, while the RP was preferentially enriched from *RPN11-TEV-ProA* cells, when purified in the absence of ATP combined with high salt washes, while the entire 26S particle (along with Blm10 and Ecm29) was enriched in the presence of ATP (Figure 6B). Strikingly, when we immunoblotted these samples for Ub, those enriched for the RP (+MG132) had a strong Ub signal, while those enriched for the CP had a weak signal (Figure 6B), thus implicating the RP as the main ubiquitylation site.

### Inhibited Proteasomes are Modified with K48 and K63 Ub-Ub Chains

The smear of Ub conjugated to inhibited proteasomes suggested that they were attached through complex architectures that might include homotypic Ub polymers as well as heterotypic Ub polymers assembled with mixed or branched Ub-Ub linkages (French et al., 2021; Haakonsen and Rape, 2019). To examine this possibility, we analyzed ubiquitylated proteasomes using linkage-specific DUBs, antibodies that recognize specific Ub-Ub connections, and yeast strains that replaced the Ub pool with K-to-R substitutions that prevent ubiquitylation at each of the seven Ub lysines. As shown in Figure 7A, when proteasome preparations (from cells treated with or without MG132) were exposed to a panel of K11, K48 and K63 linkage-specific DUBs (Hospenthal et al., 2015; Mevissen et al., 2013), whose specificity was verified by assays with synthetic dimers (Figure S7B), evidence for K48 and K63 linkages was observed. Whereas the ubiquitylation profile of inhibited proteasomes was substantially reduced when treated with a non-specific DUB USP2, no effect was seen for the K11 linkage-specific DUB Cezanne (Figure 7A). However, when treated with either the K48 or K63 linkage-specific DUBs OTUB1 and AMSH, respectively, strong Ub release was detected, implying that much of the Ub signal represented such Ub-Ub connections (Figure 7A).

**Figure 7.**
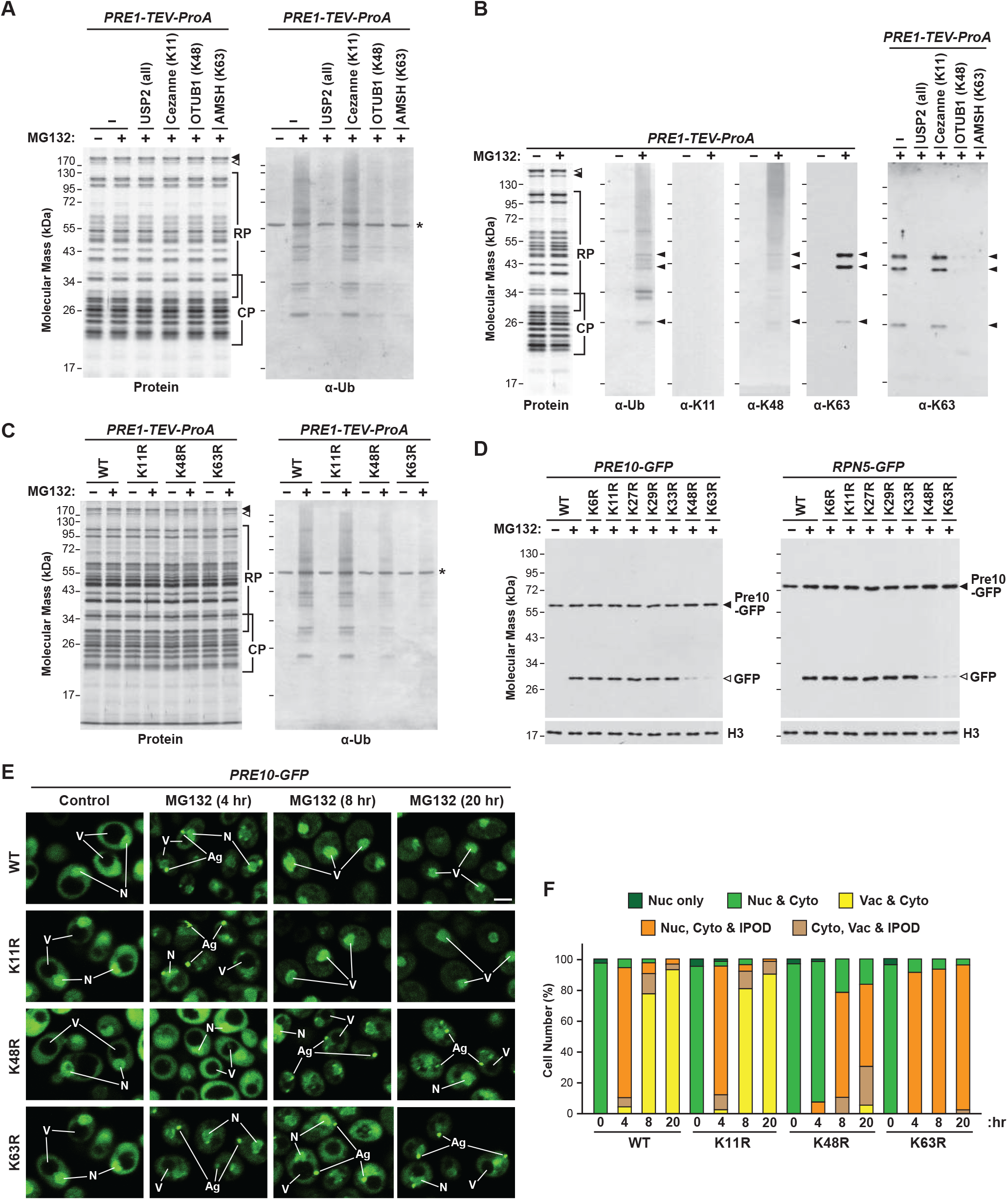
Ubiquitylation and Autophagic Transport of Inhibited Proteasomes Involves K48- and K63-Linked Poly-Ub. **(A)** Ub moieties bound to inhibited proteasomes are selectively released by DUBs specific for K48 and K63 Ub-Ub linkages. Proteasomes affinity purified from *PRE1-TEV-ProA* cells grown for 8 hr at 30°C with or without 80 µM MG132 were incubated for 30 min with the non-specific DUB USP2, or the Cezanne, OTUB1 and AMSH DUBs that are specific for K11, K48 and K63 poly-Ub linkages, respectively. Samples were subjected to SDS-PAGE, and either stained for protein or immunoblotted for Ub, as in Figure 1E. **(B)** Ubiquitylated proteasomes are specifically detected with antibodies against K48 and K63 Ub-Ub linkages. Affinity-purified proteasomes were either untreated or digested with USP2, Cezanne, OTUB1 or AMSH as in (A), and immunoblotted with anti-Ub antibodies or antibodies that specifically recognize K11, K48, and K63 Ub-Ub linkages. The arrowheads locate immunoreactive species detected by all three antibodies. **(C)** Ubiquitylation of inhibited proteasomes is suppressed in K48R and K63R cells. Proteasomes affinity purified from *PRE1-TEV-ProA* cells either wild type for Ub or expressing the K11R, K48R, and K63R Ub substitutions were analyzed as in (A). **(D)** Proteaphagy of inhibited proteasomes requires assembly of K48 and K63 Ub-Ub linkages. *PRE10-GFP* and *RPN5-GFP* cells either wild type for Ub, or where all, or nearly all (K48R), Ub genes were individually replaced with those encoding K-to-R substitutions at each of the seven Ub lysine residues, were grown for 8 hr at 30°C with or without MG132, and then assayed for free GFP release by immunoblot analysis with anti-GFP antibodies, as in Figure 1A. **(E)** Autophagic transport of inhibited proteasomes requires K48 and K63 Ub-Ub linkages. *PRE10-GFP* cells either wild type for Ub or expressing the K11R, K48R or K63R Ub substitutions were imaged by confocal fluorescence microscopy at 0, 4 or 8 hr after treatment with 80 µM MG132, as in Figure 2A. **(F)** Quantification of the cellular distribution of proteasomes in (E). Each bar represents the analysis of at least 200 cells.

Next, we immunoblotted proteasomes with Ub chain-specific antibodies (Matsumoto et al., 2010; Newton et al., 2008) whose specificity was verified with synthetic Ub dimers (Figure S7A). Whereas K11-linkage antibodies failed to recognize ubiquitylated proteasomes after MG132 treatment, those specific for K48 and K63 linkages succeeded (Figure 7B). A smear of conjugates was detected with anti-K48 linkage antibodies, while several lower molecular mass species were prominent when detected with the anti-K63 linkage antibodies, which were also modestly evident in the immunoblots with anti-K48 linkage antibodies as well as with antibodies against Ub (Figure 7B). These discrete species could be released with USP2, OUTB1 and AMSH, but not with Cezanne (Figure 7B), demonstrating not only that these species represented isopeptide-linked Ub chains, but also implying that they included both K48 and K63 Ub-Ub linkages.

Lastly, we affinity purified 26S particles (with or without MG132 treatment) from yeast strains designed to individually replace K11, K48 and K63 with arginine, thus preventing formation of Ub-Ub linkages through those sites (Meza-Gutierrez et al., 2018). Whereas the K11R and K63R strains replaced all Ub coding regions with K-to-R variants, the K48R strain replaced only ∼80% of wild-type Ub, due to the essential nature of K48-linked chains (Meza-Gutierrez et al., 2018). When inhibited proteasomes were affinity purified from these strains using the Pre1-TEV-ProA tag, only the K11R strain retained robust proteasome ubiquitylation, while those for the K48R and K63R strains were substantially suppressed (Figure 7C). Taken together, we concluded that dysfunctional proteasomes are extensively modified with Ub polymers assembled with K48- and K63-linkages.

### K48 and K63 Ub polymers Are Essential for Inhibitor-Induced Proteaphagy

Given that assembly of K48- and K63-linked Ub chains might provide a critical signal for the proteaphagy of dysfunctional particles, we tested whether yeast unable to make such polymers were also compromised in proteasome clearance. Here, we introduced the Pre10-GFP and Rpn5-GFP reporters into the complete library of K-to-R Ub substitutions (Meza-Gutierrez et al., 2018), and measured proteaphagy by the free GFP release assay and/or microscopy. When all seven K-to-R variants were assayed for free GFP after pre-treating the cells with MG132, only the K48R and K63R variants markedly slowed proteaphagy (Figure 7D). When the K11R, K48R and K63R cells harboring the Pre10-GFP reporter were then assayed by fluorescence confocal microscopy and morphometric quantification (upon treatment with or without MG132), we again saw that only the K48R and K63R variants compromised delivery of proteasomes to vacuoles (Figures 7E and 7F). Whereas proteasomes from WT and K11R cells pre-treated with MG132 exited the nucleus, aggregated into cytoplasmic aggresomes, and were delivered to vacuoles, the process was slowed in K48R cells, while for K63R cells, proteasomes successfully left the nucleus and aggregated, but stalled in subsequent vacuolar import. That both K48- and K63-linked chains were required for efficient proteaphagy upon inhibition implied that topologically complex, possibly branched, Ub chain architectures are required to enable Cue5 binding.

## DISCUSSION

Our genetic dissection of yeast proteaphagy upon proteasome inhibition revealed a surprisingly hierarchical and intricate sequence of events that translocate inactive particles from the nucleus to cytoplasmic aggresomes, and finally into vacuoles for autophagic breakdown, all of which depends on a complex set of ubiquitylation events. In fact, our studies with the *ts* E1 allele (*uba1-204*) demonstrated that Ub addition is needed early in the process with most, if not all, MG132-inhibited nuclear proteasomes remaining stuck in this compartment upon E1 inactivation. Critical steps engage three distinct E3s – San1, Rsp5 and Hul5 – along with their cognate E2s – Ubc1 and Ubc4/Ubc5 – that work collectively. Whereas the *Δsan1* mutant stalls nuclear proteasome export, *ts rsp5* and *Δhul5* mutants allow export but retain inhibited proteasomes in cytoplasmic aggresomes that appear to swell over time, implying that San1 and Rsp5/Hul5 activities are confined to the nucleus and aggresomes, respectively. Aiding this process is the oligomeric Hsp42 chaperone. While our prior studies showed that Hsp42 promotes proteasome sequestration into aggresomes (Marshall et al., 2016), we show here that Hsp42 is also required for robust 26S particle ubiquitylation, thus likely placing Hsp42 and proteasome condensation downstream of San1 but upstream of Rsp5 and Hul5. Finally, a complicated poly-Ub chain architecture is assembled mostly on the RP that includes a mix of K48 and K63 Ub-Ub linkages, which we presume provides the necessary code for Cue5 recognition.

Notably, when surveying the literature on San1, Rsp5 and Hul5, along with their associated E2s, it became apparent that the ubiquitylation cascade revealed here is not confined to degrading inactive proteasomes but likely participates in numerous, more general PQC routes, and thus could define a core cellular mechanism for maintaining proteostasis. For example, nuclear-localized San1 has been well connected to PQC in this compartment, works with the Ubc1, Ubc4 and Ubc5 E2s, and is known to catalyze assembly of K48-linked poly-Ub chains (Gardner et al., 2005; Ibarra et al., 2016; Matsuo et al., 2011; Samant et al., 2018). For proteaphagy, San1 appears essential for the nuclear export of inhibited proteasomes, possibly in concert with Ubc4/Ubc5 given that proteasomes in *Δubc4 Δubc5* cells remain nuclear, as in *Δsan1* cells, while proteasomes in *Δubc1* cells do exit nuclei, albeit more slowly.

Similarly, Rsp5 and its human ortholog Nedd4 have been associated with PQC more broadly, such as targeting cytosolic misfolded proteins that arise during heat stress (Fang et al., 2014), ribophagy (Kraft and Peter, 2008), and the autophagic clearance of aggregation-prone substrates in conjunction with Ubc4/Ubc5 and Cue5 (Lu et al., 2014; Tardiff et al., 2013; Tofaris et al., 2011). Rsp5 has been localized to ill-defined cytoplasmic puncta near the plasma membrane and adjacent to vacuoles that could represent the aggresomes seen here (Wang et al., 2001). Rsp5 specifically catalyzes the formation of K63-linked poly-Ub chains (Saeki et al., 2009), with the proteasome RP subunit Rpn10 even identified as one Rsp5 substrate (Isasa et al., 2010). Because Rsp5 and Ubc4/Ubc5 also direct proteins to the UPS, Lu et al. (2017) proposed that the soluble versus aggregated nature of the substrate determines whether it enters the UPS or autophagy after Rsp5 ubiquitylation.

Finally, Hul5 has been similarly implicated in the turnover of misfolded and low solubility proteins, with its cytoplasmic distribution influenced by proteotoxic stress (Fang et al., 2011). It also directly associates with proteasomes substoichiometrically (Leggett et al., 2002), especially particles that are structurally impaired (Park et al., 2011), where it has been proposed to provide an E4 activity that further modifies ubiquitylated substrates (Crosas et al., 2006). We hypothesize that Hul5 associates with aggresome-bound proteasomes awaiting autophagic clearance after their ubiquitylation by Rsp5.

We also emphasize that other yeast E3s connected to PQC were not found to influence proteaphagy. Included were the inner nuclear membrane quality control Asi complex (Foresti et al., 2014), scaffold proteins of SCF E3s (Cul3, Cdc53, Rtt101 and Skp1) (Finley et al., 2012), Hel2 and Ltn1 involved in ribosome-associated quality control (Brandman et al., 2012), Hrd1 and Doa10 that mediate ER-associated proteolysis (Mehrtash and Hochstrasser, 2019), and Ubr1 and Ufd4 that participate in cytoplasmic PQC and interact with proteasomes (Heck et al., 2010; Xie and Varshavsky, 2000). The Ub receptor Dsk2, which has been implicated in nuclear PQC (Samant et al., 2018), was also not required for clearing inhibited proteasomes (Marshall et al., 2016). These exclusions imply that San1, Rsp5 and Hul5 participate in a unique form of selective PQC possibly designed to remove aggregated substrates by autophagy (Lu et al., 2014).

How inactive proteasomes are first recognized as proteaphagy substrates is not yet clear, but because San1 is required early, its activity might be crucial. Given that San1 has been reported to bind intrinsically disordered regions (IDRs), possibly through its own structural disorder (Hickey et al., 2021; Rosenbaum and Gardner, 2011), and that a number of proteasomes RP subunits begin or terminate in predicted IDRs (Aufderheide et al., 2015), we speculate that changes in IDR accessibility upon proteasome inhibition are key signals. It is also possible that these same IDRs (or subsequently bound Ub moieties) in concert with Hsp42 also promote entry of inhibited proteasomes into aggresomes, given that Hsp42 oligomers can sequester misfolded/insoluble proteins (Specht et al., 2011).

Prior studies have connected robust ubiquitylation to many forms of selective autophagy, including the removal of protein aggregates via receptors such as NBR1, p62/SQSTM1, Tollip and Cue5 (Lu et al., 2014; Yin et al., 2020). Our studies with linkage-specific Ub antibodies and DUBs, and yeast strains substituting each of the seven Ub lysines, revealed an essential role for K48- and K63-linked Ub-Ub chains, but not those involving K6, K11, K27, K29, or K33, in the autophagic clearance of dysfunctional proteasomes. The failure to need K11 linkages was unexpected given the reported importance of heterotypic K11/K48-linked branched polymers during aggregation-prone protein clearance (Yau et al., 2017). While the exact topolog(ies) of proteasome-bound Ub linkages are not yet known, our data implicate heterotypic polymers assembled with mixed or branched Ub-Ub linkages as possibilities. Included is the robust release of most Ub signals from inhibited proteasomes when treated individually with the K48-specific and K63-specific DUBs, and the need for both K48 and K63 Ub-Ub linkages for robust proteaphagy. Probing inhibited proteasome preparations with antibodies against K48 and K63 Ub-Ub linkages revealed a striking difference in ubiquitylation profiles, with the K48-specific antibodies mainly detecting a smear of high molecular mass conjugates, while the K63-specific antibodies only detected a few prominent lower molecular mass species. These discrete species were sensitive to OTUB1 and AMSH, implying that they contain both K48 and K63 Ub-Ub linkages, and thus could represent the core Ub polymer architecture based on their low apparent molecular mass. Clearly, further linkage analysis of these species by advanced mass spectrometric methods (French et al., 2021; Haakonsen and Rape, 2019), along with the identification of the modified proteasome subunit(s), should be revealing.

How the proteaphagy machinery generates such a complex ubiquitylation pattern upon proteasome inhibition is unclear, but a model is possible based on the activities/locations of the components involved. We propose that San1, along with Ubc1 and Ubc4/Ubc5, recognize inactive proteasomes and either monoubiquitylate the particles or assemble poly-Ub species through Ubc1 that can build K48-linked chains (Pluska et al., 2021). After nuclear export and condensation with the help of Hsp42, aggresome-bound proteasomes are further modified with K63-linked Ub polymers by Rsp5. Hul5 then assembles mixed or branched K48-linked Ub chains onto existing K63 Ub polymers, with HECT E3s such as Rsp5 and Hul5 being uniquely designed for such chain extensions through their formation of E3-Ub adducts prior to Ub transfer (French et al., 2021). This complex Ub code is then uniquely recognized by Cue5 through its CUE Ub-binding surface, which can bind both K48 and K63-linked Ub chains (Lu et al., 2014).

Such a synchronized scheme for E3s is not without precedent, but has not yet been described for autophagy. As examples, yeast Ufd2 and Ufd4 work together to synthesize branched K29/K48 chains on substrates (Liu et al., 2017), while in humans, branched K48/K63 chains are produced by TRAF6 and HUWE1 during NF-κB signaling, and by ITCH and UBR5 during apoptosis (Ohtake et al., 2016; 2018). One likely advantage of using E3s with distinct chain preferences is to spatially and/or temporally separate ubiquitylation marks with different consequences. For illustration, the pro-apoptotic regulator TXNIP is first modified with non-proteolytic K63-linked chains by ITCH, before UBR5 attaches K48-linked Ub moieties to generate branched K48/K63 chains that target TXNIP for proteasomal degradation (Ohtake et al., 2018).

In conclusion, our studies on the events underpinning the proteaphagy of dysfunctional proteasomes identified Hsp42 and a hierarchical trio of E3s and their associated E2s that sequentially drive nuclear export, proteasome aggregation, and autophagic clearance, ultimately through assembly of a topologically complex Ub code containing both K48 and K63 Ub-Ub linkages. As the central components San1, Rsp5, Hul5, Hsp42, and Cue5 also participate in proteostasis more generally (Lu et al., 2014; Specht et al., 2011), proteaphagy likely engages a common route for eliminating aberrant proteins, and thus represents an tractable model for understanding PQC checkpoints that remove amyloidogenic proteins, including those associated with aggregation-prone pathologies.

**Table 1.** *Saccharomyces cerevisiae* Strains Used in This Study; Related to STAR Methods.

**Table 2.** Accession Numbers of Genes and Proteins Used in This Study; Related to STAR Methods.

**Table 3.** Oligonucleotide Primers Used in This Study; Related to STAR Methods.

## ACKNOWLEDGEMENTS

We wish to thank E. Sethe Burgie, Pedro Carvalho, Raymond J. Deshaies, Daniel Finley, Audrey P. Gasch, Richard G. Gardner, Christopher M. Hickey, Mark Hochstrasser, Daniel J. Klionsky and Heather L. True for generously sharing reagents and strains, and Dianne M. Duncan, Sujina Mali and Jerry Wei for technical support. This work was supported by a grant from the National Institutes of Health, National Institute of General Medical Sciences (R01-GM124452) to R.D.V.

## AUTHOR CONTRIBUTIONS

R.S.M. and R.D.V. conceived the project and experimental approaches, R.S.M. performed the research, and R.S.M. and R.D.V. analyzed the data and wrote the paper.

## STAR METHODS

### RESOURCE AVAILIBILITY

#### Lead Contact

Further information and requests for resources and reagents should be directed to and will be fulfilled by the lead contact: Richard D. Vierstra.

#### Materials Availability

All unique materials generated during this study will be made available upon request.

#### Data and Code Availability

This study did not generate or analyze datasets or code.

### EXPERIMENTAL MODEL AND SUBJECT DETAILS

#### Yeast Strains and Manipulations

Unless otherwise stated, all yeast manipulations were performed according to standard protocols (Dunham et al., 2015; Marshall et al., 2016). Details of all strains used in this study are provided in Table S1, and all relevant *Saccharomyces* Genome Database identifiers are listed in Table S2. Strains expressing Pre10-GFP and Rpn5-GFP in the BY4741 background (Brachmann et al., 1998; Marshall et al., 2016) were obtained from the yeast GFP clone collection (Life Technologies) and cultured on synthetic dropout medium lacking histidine. Deletion strains in the BY4742 background were generously provided by Mark Hochstrasser (Yale University) from the TUS2.0 collection (Hickey et al., 2021) and were cultured on YPDA (+N) medium containing 200 µg/ml Geneticin, except for the *Δerg6* deletion (Marshall et al., 2016), which was grown on YPDA medium containing 200 µg/ml hygromycin B. The yeast strain collection harboring K-to-R substitution mutations for the seven Ub lysines in the SK1 background was as described (Mez-Gutierrez et al., 2018). All other strains were generously provided by collaborators, including from the yeast *ts* conditional mutant collection (Li et al., 2011).

For time course experiments, 15-ml liquid cultures in YPDA (+N) medium were grown overnight at 30°C with vigorous shaking, diluted to an OD_600_ of 0.1 in 15 ml, grown for an additional 2-3 hours until an OD_600_ of approximately 0.5 was reached, transferred to 37°C for 1 hr if required, switched to +N medium, –N medium, or +N medium containing 80 µM MG132 ((*N*-benzyloxycarbonyl)-leucinyl-leucinyl-leucinal), and then grown at 30°C or 37°C for a further 8 hr. For –N treatment, cultures were re-suspended in medium lacking N (0.17% (w/v) yeast nitrogen base without amino acids and ammonium sulphate, 2% (w/v) glucose). All experiments involving MG132 exploited the *Δerg6* deletion that eliminated ergosterol biosynthesis to enhanced membrane permeability (Lee and Goldberg, 1996). Cell aliquots corresponding to 1.5 OD units were collected by centrifugation, washed, and immediately frozen in liquid nitrogen.

### METHOD DETAILS

#### Screening the Targeted Ubiquitin System (TUS) Deletion Library

For screens of the TUS library, *PRE10-GFP* and *RPN5-GFP* strains (mating type α) were mated with each TUS mutant (mating type a). For *ts* or proteasome purification strains of mating type α, mating type switching was performed using standard protocols following transformation with the pGAL::HO plasmid (a gift from Heather L. True, Washington University in St. Louis). Cells were streaked together in a patch onto solid YPDA medium and grown for 2 days at 30°C. The resulting cells (and their individual parent strains) were then streaked out onto double selective medium (synthetic dropout medium lacking histidine and containing 200 µg/ml Geneticin), and grown for a further 2 days at 30°C. The resulting colonies were presumed to be diploids if neither parental strain grew on the double selection. To induce sporulation, diploid cells were patched onto freshly made GNA pre-sporulation plates (1% (w/v) yeast extract, 1.125% (w/v) beef extract, 1.875% (w/v) bacto-peptone, 5% (w/v) glucose, 2% (w/v) bacto-agar), grown overnight at 30°C, then re-patched onto GNA plates and grown overnight for a second time. Colonies were suspended in 2 ml of liquid sporulation medium (1% (w/v) potassium acetate, 0.005% (w/v) zinc acetate), and aliquots of 100 µl were transferred into a 96-well plate and incubated with mixing for 5 days at 25°C, followed by 3 days at 30°C.

To isolate spores from the resulting ascus, cells were collected by centrifugation and re-suspended in 50 µl water containing 50 U lyticase and 0.5 µl β-mercaptoethanol. Cells were incubated overnight at 30°C with gentle shaking, 50 µl of 1.5% (v/v) Triton X-100 was added, and cells were incubated on ice for 15 min. The plate was then tightly sealed and disrupted by thrice-repeated sonication in a water bath (Digital Pro Ultrasonic). The spores were washed twice in 100 µl of 1.5% (v/v) Triton X-100, sonicated again as above, washed twice in H_2_O, and resuspended in H_2_O at ∼1,000 spores per ml. Aliquots (200 µl) were plated onto double selective medium and grown at 30°C for three days. The resulting colonies were confirmed as haploid and containing the desired deletions by PCR genotyping using primer pairs listed in Table S3. Expression of the GFP fusions was confirmed by fluorescence confocal microscopy.

#### Gene Cloning and Site-directed Mutagenesis

To clone the E1, E2 and E3 coding sequences, total yeast RNA was first extracted from log-phase BY4741 cells using the RNeasy mini kit (QIAGEN), and then converted into cDNA using the SuperScript III first-strand synthesis system (Thermo Fisher Scientific) and oligo(dT)_20_ primers. Coding sequences amplified by PCR were recombined into pDONR221 via the Gateway BP clonase II reaction (Thermo Fisher Scientific), altered by site-directed mutagenesis using the QuikChange II site-directed mutagenesis kit (Agilent Technologies) as needed, sequence-verified, and recombined in-frame with the pAG423-GPD-ccdB-DsRed, pAG424GPD-ccdB-HA, pAG425-GPD-ccdB-EGFP, pAG425-GPD-EGFP-ccdB or pAG425-GPD-ccdB vectors (Addgene, product numbers 14366, 14248, 14202, 14322 and 14154, respectively) via the Gateway LR clonase II reaction (Thermo Fisher Scientific). The sequences encoding N-terminal HA-tags (YPYDVPDYA) were incorporated into the appropriate PCR amplification primers. Plasmid transformation was performed by the lithium acetate procedure (Dunham et al., 2015), and the transformed cells were cultured on synthetic dropout medium lacking leucine, histidine and/or tryptophan as required.

#### Immunological Techniques

For analysis of total protein extracts, frozen cell aliquots were re-suspended in 500 µl of lysis buffer (0.2 N NaOH and 1% (v/v) β-mercaptoethanol), followed by precipitation with 50 µl of 50% (w/v) trichloroacetic acid. The precipitates were collected by centrifugation at 4°C, washed once with 1 ml of ice-cold acetone, re-suspended in 150 µl of SDS-PAGE sample buffer, and heated at 95°C for 5 min (Marshall et al., 2016). For affinity-purified proteasomes, the samples were directly solubilized in SDS-PAGE sample buffer as above. After SDS-PAGE, the gels were stained for protein with silver or subjected to immunoblot analysis using Immobilon-P PVDF membranes (Milllipore), as previously described (Marshall et al., 2016). The anti-K11, anti-K48 and anti-K63 Ub antibodies were also as described (Matsumoto et al., 2010; Newton et al., 2008). Details of all primary and secondary antibodies used are given in the Key Resources Table. Immunoblots with anti-Ub or anti-K48 antibodies were developed with the 1-Step NBT/BCIP AP Substrate kit (Thermo Fisher Scientific); others were developed using the SuperSignal West Pico Plus Chemiluminescent Substrate or the SuperSignal West Femto Maximum Sensitivity Substrate (both from Thermo Fisher Scientific).

#### Proteasome Affinity Purifications

Affinity purification of proteasomes via the CP or RP was performed as previously described (Leggett et al., 2002; Marshall et al., 2016). Yeast strains in which the Pre1 or Rpn11 subunits had been genetically replaced by variants tagged with TEV-cleavable Protein A (ProA) (Leggett et al., 2002) were grown as above at 30°C or 37°C in +N medium with or without 80 µM MG132. Briefly, frozen cell pellets were ground to a fine powder and rehydrated on ice with 1 volume of lysis buffer (50 mM Tris-HCl (pH 7.5), 5 mM MgCl_2_, 1 mM EDTA, 10% (v/v) glycerol, with 2 mM ATP (unless otherwise indicated), 2 mM PMSF, 1 mM benzamidine, 10 µg/ml pepstatin A, 1 µg/ml antipain and 1X protease inhibitor cocktail (Sigma-Aldrich) added immediately before use). Filtered and clarified extracts were affinity purified with 100 µl of whole molecule IgG beads (MP Biomedicals) pre-equilibrated in lysis buffer, washed three times with 2 ml of proteasome wash buffer (50 mM Tris-HCl (pH 7.5), either 50 or 250 mM NaCl, 5 mM MgCl_2_, 1 mM EDTA, 2 mM ATP (unless otherwise indicated), 10% (v/v) glycerol), and twice with 1 ml of TEV protease buffer (50 mM Tris-HCl (pH 7.5), 5 mM MgCl_2_, 1 mM EDTA, 2 mM ATP, 1 mM DTT, 10% (v/v) glycerol). Bound proteins were eluted by incubating the beads for 1 hr at 30°C with 300 µl of TEV protease buffer containing 20 ng/µl of recombinant 6His-TEV (a gift from E. Sethe Burgie, Washington University in St. Louis). Where indicated, DUB treatments were performed prior to elution (see below). Residual 6His-TEV was removed by subsequent incubation of the eluates with 50 µl nickel-nitrilotriacetic acid (Ni-NTA)-agarose beads (QIAGEN) pre-equilibrated in TEV protease buffer containing 40 mM imidazole (for a final concentration of 10 mM).

#### DUB Treatments

The USP2, Cezanne, OTUB1 and AMSH DUBs were purchased from Boston Biochem (catalog numbers E-504, E-563-050, E-522B-050 and E-4548B, respectively). Where indicated, IgG beads decorated with affinity-purified proteasomes were incubated prior to elution for 30 min at 30°C with each DUB at a concentration of 10 nM in a volume of 300 µl. The beads were washed three times with 1 ml of TEV protease buffer to remove the DUB and released Ub, before elution of the proteasomes by TEV protease cleavage as above. For confirmation of the reported cleavage specificities, the DUBs were incubated at 30°C for various times with 25 µg of pure Ub dimers isopeptide-linked via K11, K48 or K63 linkages (Boston Biochem, catalog numbers UC-40B-025, UC-200B-025 and UC-300B, respectively) resuspended in 50 mM Tris-HCl (pH 7.5), 50 mM NaCl, and 10 mM DTT (Hospenthal et al., 2015; Mevissen et al., 2013). Samples were analysed for digestion by silver staining following SDS-PAGE.

#### Confocal Microscopy

Cells expressing Pre10-GFP were grown in +N or –N media with or without MG132 as described above, and visualized by fluorescence confocal microscopy with a Nikon A1^+^ microscope using a 100X oil objective (numerical aperture 1.46). GFP excitation was performed at 488 nm, and emission was collected between 500 and 530 nm. Cells were fixed using cover slips coated with a 2 mg/ml solution of concanavalin A and then air dried (Marshall et al., 2016). To minimize auto-fluorescence from the YPDA medium, cells were resuspended in synthetic dropout medium lacking appropriate amino acids prior to imaging. All confocal images were scanned in single track mode and processed using Elements Viewer (Nikon Imaging Software) and/or Adobe Photoshop CC, before conversion to TIFF files for use in the figures. Within each figure, all images were captured using identical microscope settings.

#### Pho8Δ60 Enzyme Assays

The Pho8Δ60 assays were performed as previously described (Noda and Klionsky, 2008), with minor modifications (Marshall et al., 2016). Strains TN124 or RSM879-892 were grown in 250 ml cultures at 30°C or 37°C, transferred to –N medium, and aliquots corresponding to 5.0 OD units were collected by centrifugation at the indicated time points. Clarified lysates were analysed for Pho8Δ60 activity at 37°C in 250 mM Tris-HCl (pH 8.5), 10 mM MgSO_4_, 10 µM ZnSO_4_, 1% (v/v) Triton X-100 and 1.5 mM *p*-nitrophenyl phosphate (Sigma-Aldrich), and quantified by the absorbance of the *p*-nitrophenol product at 400 nm.

### QUANTIFICATION AND STATISTICAL ANALYSES

#### Statistical analysis

Datasets were statistically analysed by one-way analysis of variance (ANOVA) to determine the presence or absence of significant differences, followed by Tukey’s post-hoc tests to identify data points that were significantly different from one another.

### SUPPLEMENTAL INFORMATION

Supplemental Information includes 7 figures, 3 tables, and can be found with this article online at xxx.

## SUPPLEMENTAL FIGURE LEGENDS

**Figure S1.**
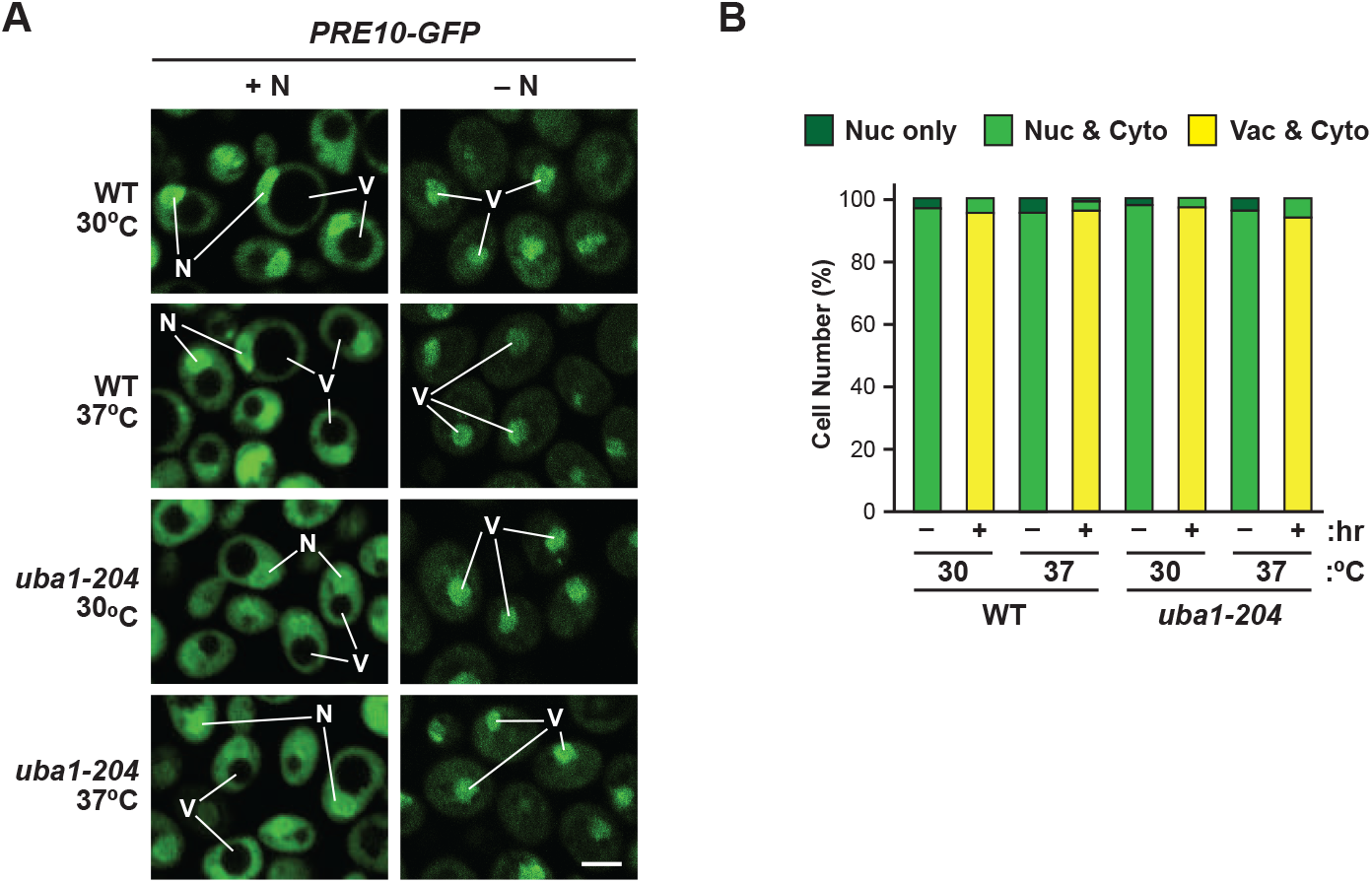
Ubiquitylation is Not Essential for Delivering Yeast Proteasomes to the Vacuole by Autophagy During Nitrogen Starvation. Related to Figure 1. **(A)** Proteasomes are transported to the vacuole in WT and *uba1-204* cells upon N starvation at the non-permissive temperature. WT and mutant *PRE10-GFP* cells were grown on +N medium at 30°C and either kept on +N medium or switched to –N medium, grown for 8 hr at either 30°C or 37°C, and then imaged by confocal fluorescence microscopy. N, nucleus; V, vacuole; Ag, cytoplasmic aggregate. Scale bar, 2 µm. **(B)** Quantification of the cellular distribution of proteasomes as visualized in (A). Each bar represents the analysis of at least 200 cells.

**Figure S2.**
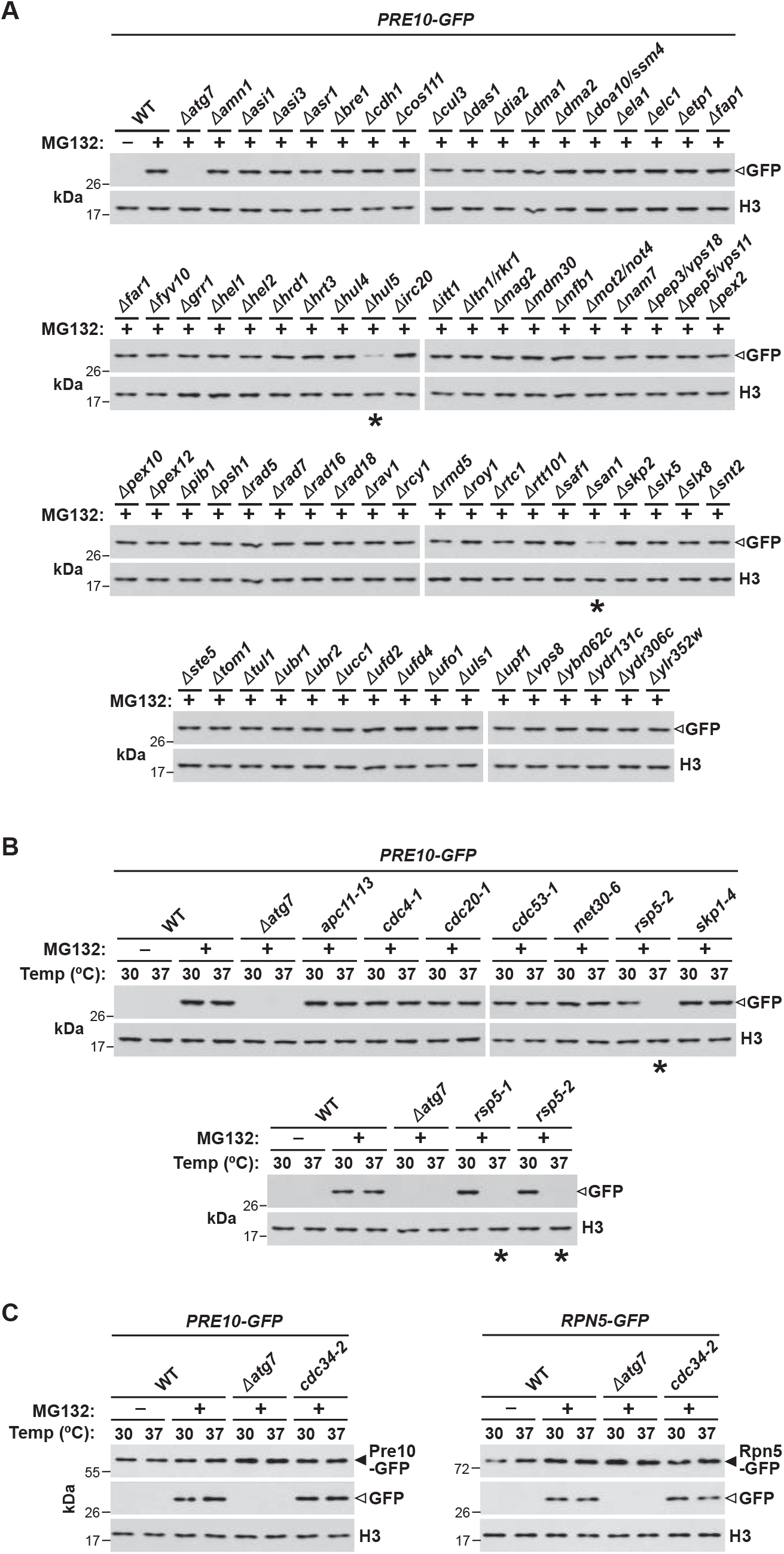
Full Screen of Available Yeast E2 and E3 Mutants for Their Impact on Proteaphagy of Inhibited Proteasomes. Related to Figures 3 and 5. **(A)** A screen of 81 available deletion mutants of non-essential yeast E3s in the TUS collection identified Hul5 and San1 as important for proteaphagy of inhibited proteasomes. Cells expressing *PRE10-GFP* were grown for 8 hr at 30°C on +N medium with or without 80 µM MG132 and assayed for the release of free GFP from the reporters by immunoblot analysis of cell extracts with anti-GFP antibodies. **(B)** A screen of *ts* alleles impacting seven essential yeast E3s or E3 components identified Rsp5 as important for proteaphagy of inhibited proteasomes. WT and mutant *PRE10-GFP* cells were grown for 8 hr at either 30°C or 37°C with or without MG132 and then assayed for the release of free GFP as in (A). **(C)** The *ts cdc34-2* allele of the Cdc34/Ubc3 E2 does not block proteaphagy of inhibited proteasomes at the non-permissive temperature. WT and *cdc34-2* cells expressing *PRE10-GFP* (left panel) or *RPN5-GFP* (right panel) were grown for 8 hr at either 30°C or 37°C with or without MG132 and then assayed for the release of free GFP as in (A). For panels (A and B), only sections of the gels harboring free GFP (open arrowhead) are shown. Immunodetection of histone H3 was included to confirm near equal protein loading. The *Δatg7* mutant was included as a positive control. Asterisks highlight the *Δhul5*, *Δsan1*, *Δhul5 Δsan1*, *rsp5-1* and *rsp5-2* strains. See Figures 3A, 3B, and 5A for confirmation of the results.

**Figure S3.**
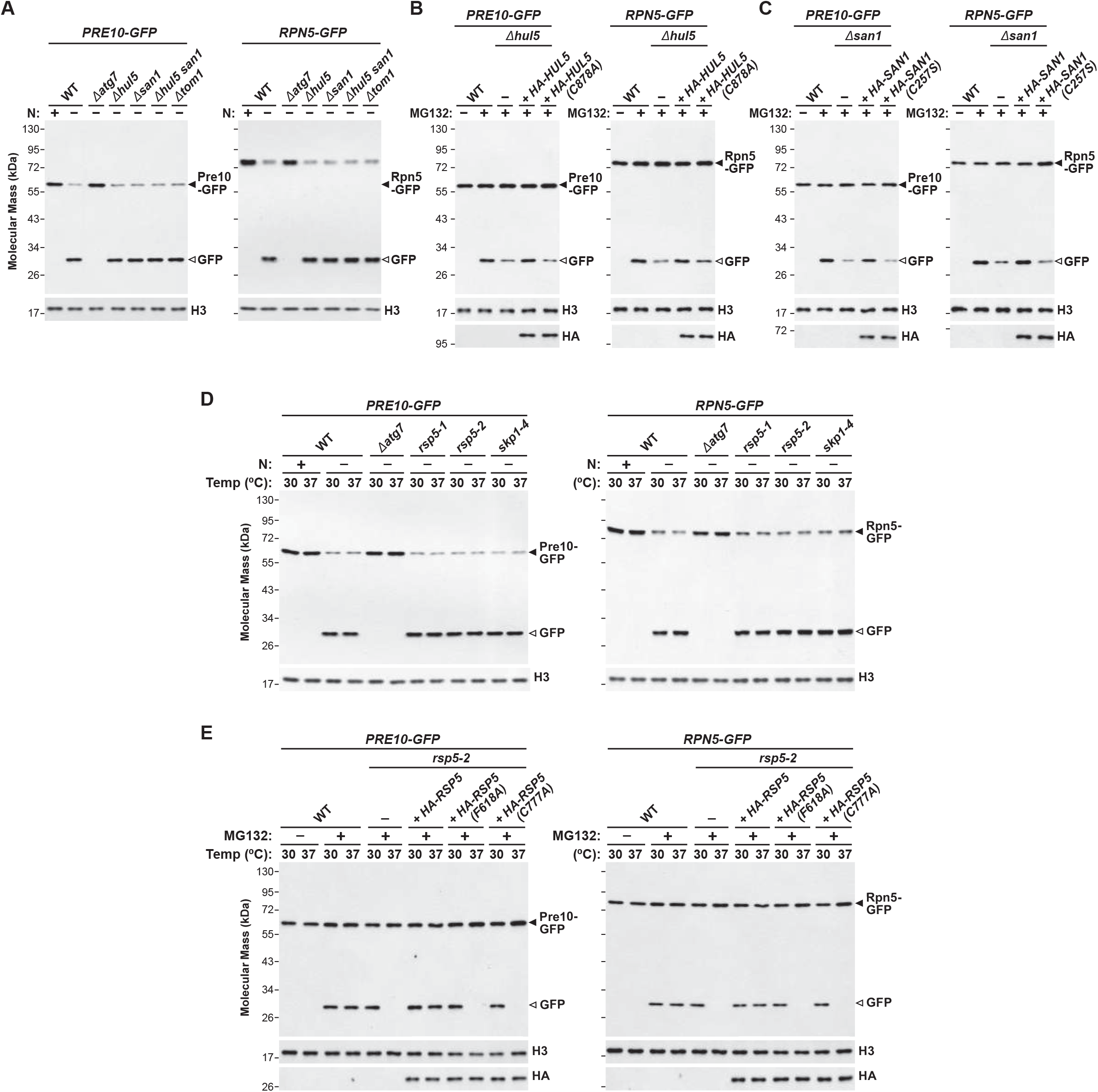
Full Description of the Impacts of Hul5, Rsp5, and San1 E3s on the Autophagic Degradation of Proteasomes. Related to Figure 3. **(A)** Proteaphagy proceeds normally in *Δhul5* and *Δsan1* cells during N starvation. WT and *Δhul5, Δsan1* and *Δhul5 Δsan1* cells expressing *PRE10-GFP* (left panel) or *RPN5-GFP* (right panel) were grown in +N medium at 30°C and either kept in +N medium or switched to –N medium and grown for an additional 8 hr. Cells were assayed for the release of free GFP from the reporters by immunoblot analysis of total cell extracts with anti-GFP antibodies. Immunodetection of histone H3 was included to confirm near equal protein loading. Closed and open arrowheads locate the GFP fusions and free GFP, respectively. The *Δatg7* and *Δtom1* mutants were included as positive and negative controls, respectively. **(B)** Proteaphagy of inhibited proteasomes requires enzymatically active Hul5. WT and *Δhul5* cells expressing *PRE10-GFP* (left panels) or *RPN5-GFP* (right panels), either alone or complemented with HA-tagged versions of wild-type Hul5 or the active site mutant Hul5(C878A), were grown for 8 hr in +N medium at 30°C with or without 80 µM MG132. Cells were assayed for the release of free-GFP from the reporters by immunoblot analysis as in (A). Immunoblot analysis with anti-HA antibodies confirmed expression of the HA-tagged Hul5 variants. **(C)** Proteaphagy of inhibited proteasomes requires enzymatically active San1. WT and *Δsan1* cells expressing *PRE10-GFP* (left panels) or *RPN5-GFP* (right panels), either alone or complemented with HA-tagged versions of wild-type San1 or the active site mutant San1(C257S), were grown for 8 hr in +N medium at 30°C with or without MG132 and assayed for the release of free GFP by immunoblot analysis as in (B). Immunoblot analysis with anti-HA antibodies confirmed expression of the HA-tagged San1 variants. **(D)** Proteaphagy proceeds normally in *ts rsp5* cells during N starvation at the non-permissive temperature. WT, *rsp5-1* and *rsp5-2* cells expressing *PRE10-GFP* (left panel) or *RPN5-GFP* (right panel) were grown in +N medium at 30°C and either kept in +N medium or switched to –N medium and grown for an additional 8 hr at 30°C or 37°C. Cells were assayed for the release of free GFP from the reporters by immunoblot analysis as in (A). The *Δatg7* and *ts skp1-4* mutants were included as positive and negative controls, respectively. **(E)** Proteaphagy of inhibited proteasomes requires active Rsp5. WT and *rsp5-2* cells expressing *PRE10-GFP* (left panels) or *RPN5-GFP* (right panels), either alone or complemented with HA-tagged versions of wild-type Rsp5 or the Ub-binding Rsp5(F618A) or active site Rsp5(C777A) mutants, were grown for 8 hr in +N medium at 30°C or 37°C with or without MG132. Cells were assayed for the release of free-GFP from the reporters by immunoblot analysis as in (A). Immunoblot analysis with anti-HA antibodies confirmed expression of the HA-tagged Rsp5 variants.

**Figure S4.**
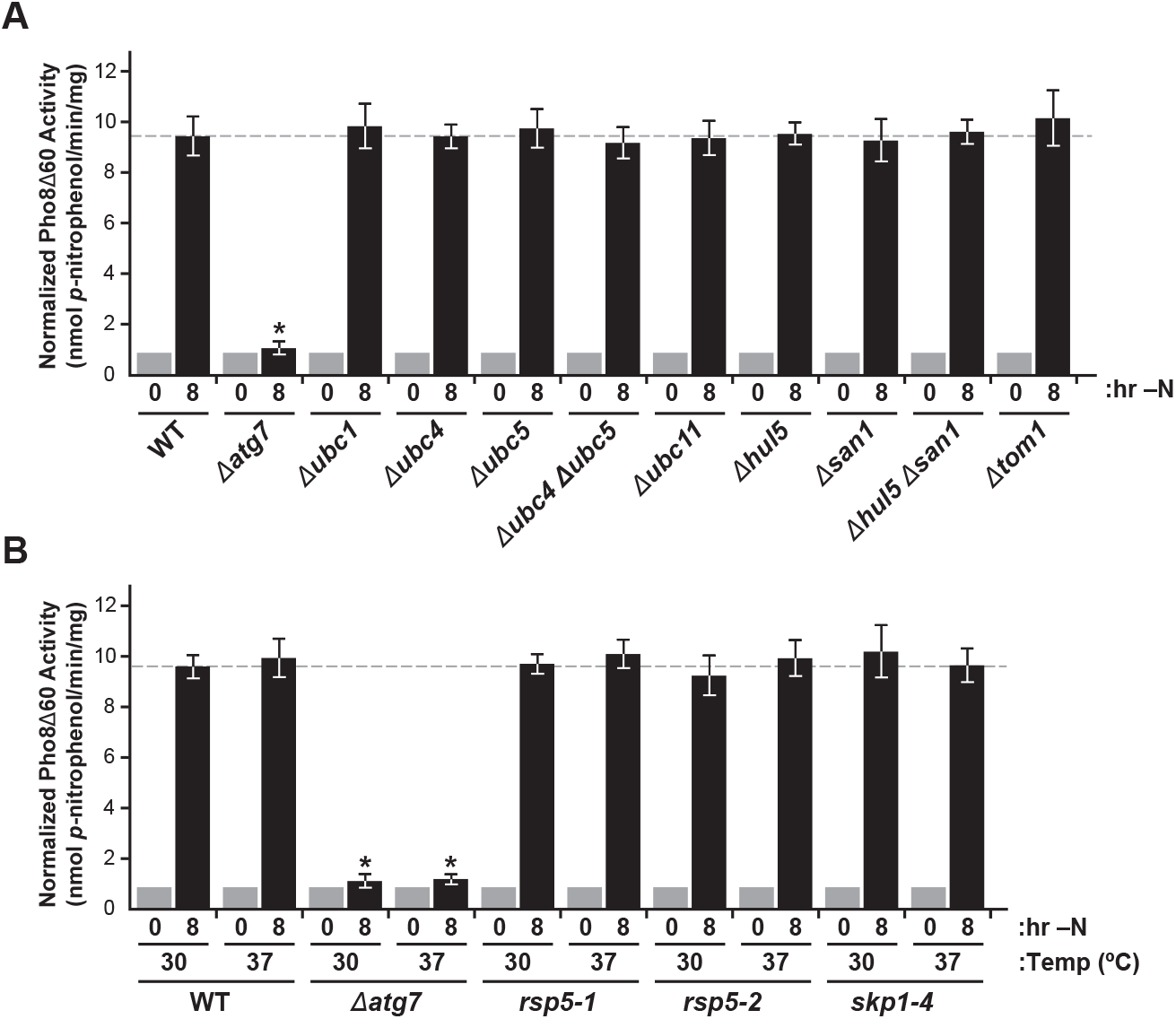
Loss of the Uba1 E1, the Ubc1, Ubc4 and Ubc5 E2s, and the Hul5, Rsp5 and San1 E3s does not Compromise Bulk Autophagy Induced by N Starvation. Related to Figures 4 and 5. Bulk autophagy was assayed by the Pho8Δ60 reporter, whose phosphatase activity depends on its autophagic transport and subsequent vacuolar processing. Each bar represents the mean (±SD) of three biological replicates, each with three technical replicates, measuring Pho8Δ60 activity at time 0 and 8 hr after switching cells grown in +N medium at 30°C to –N medium and grown at 30°C or 37°C. The dashed line reflects the response of WT. The *Δatg7* and the *Δubc11*, *Δtom1* and *skp1-4* mutants were included as positive and negative controls, respectively. **(A)** Analysis of the deletion mutants *Δubc1*, *Δubc4*, *Δubc5*, *Δubc4 Δubc5*, *Δhul5*, *Δsan1* and *Δhul5 Δsan1*. **(B)** Analysis of the *ts uba1-204*, *rsp5-1* and *rsp5-2* alleles.

**Figure S5.**
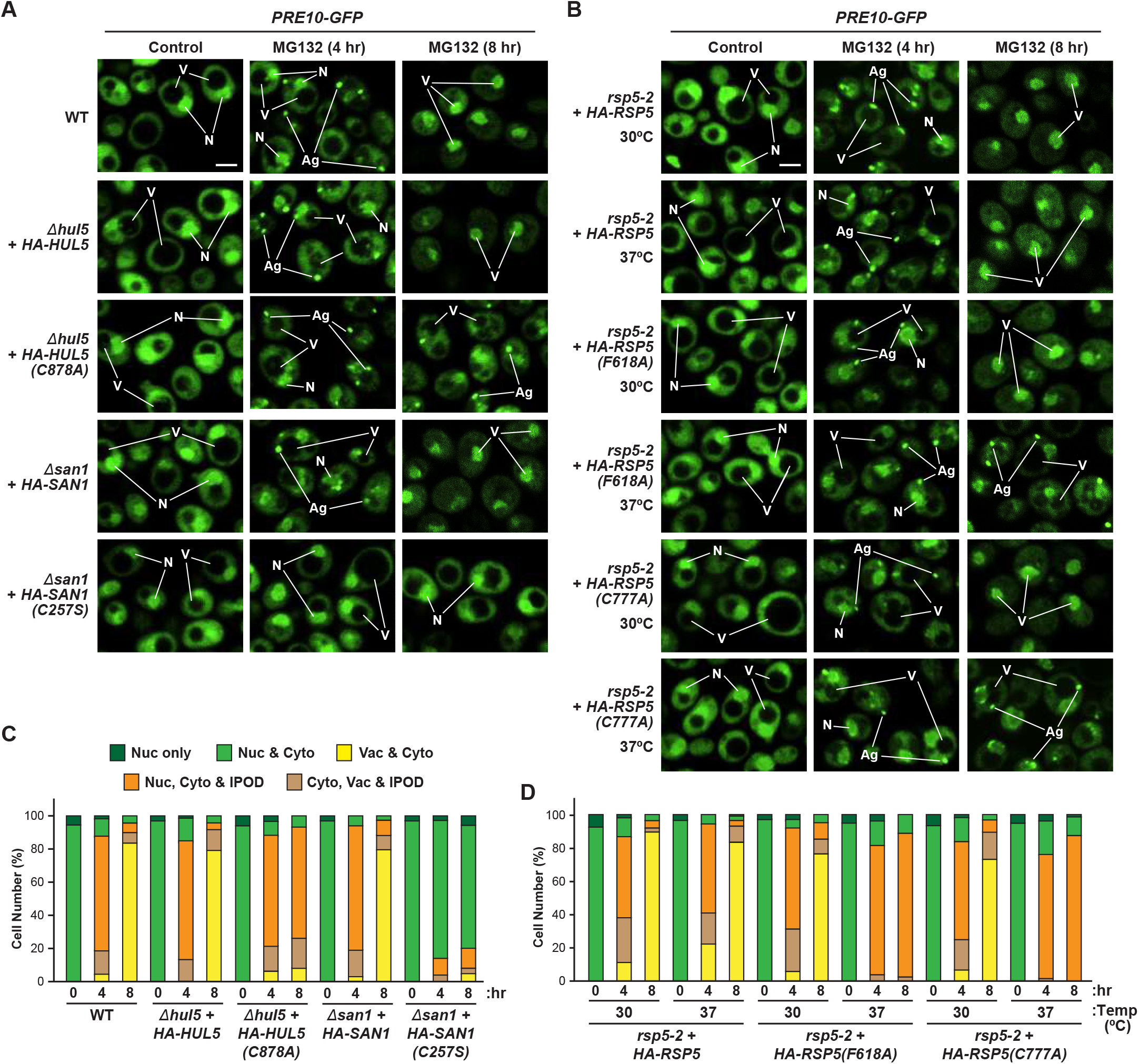
Effects of the *Δhul5*, *Δsan1* and *rsp5-2* Mutations on the Delivery of Inhibited Proteasomes to the Vacuole During Proteaphagy. Related to Figure 4. **(A)** Inhibited proteasomes are transported to vacuoles in WT and in *Δhul5* and *Δsan1* cells complemented with wild-type Hul5 or San1, respectively, but not with the Hul5(C878A) or San1(C257S) active site mutants. *PRE10-GFP* cells were grown at 30°C in +N medium and imaged by confocal fluorescence microscopy at 0, 4 or 8 hr after treatment with 80 µM MG132. N, nucleus; V, vacuole; Ag, cytoplasmic aggregate. Scale bar, 2 µm. **B)** Inhibited proteasomes are transported to vacuoles in WT and in *rps5-2* cells complemented with wild-type Rsp5, but not with the Rsp5(F618A) or Rsp5(C777A) mutants. *PRE10-GFP* cells were grown at 30°C or 37°C in +N medium and imaged by confocal fluorescence microscopy at 0, 4 or 8 hr after MG132 treatment as in (A). **(C** and **D)** Quantification of the cellular distribution of proteasomes in (A) and (B), respectively. Each bar represents the analysis of at least 200 cells.

**Figure S6.**
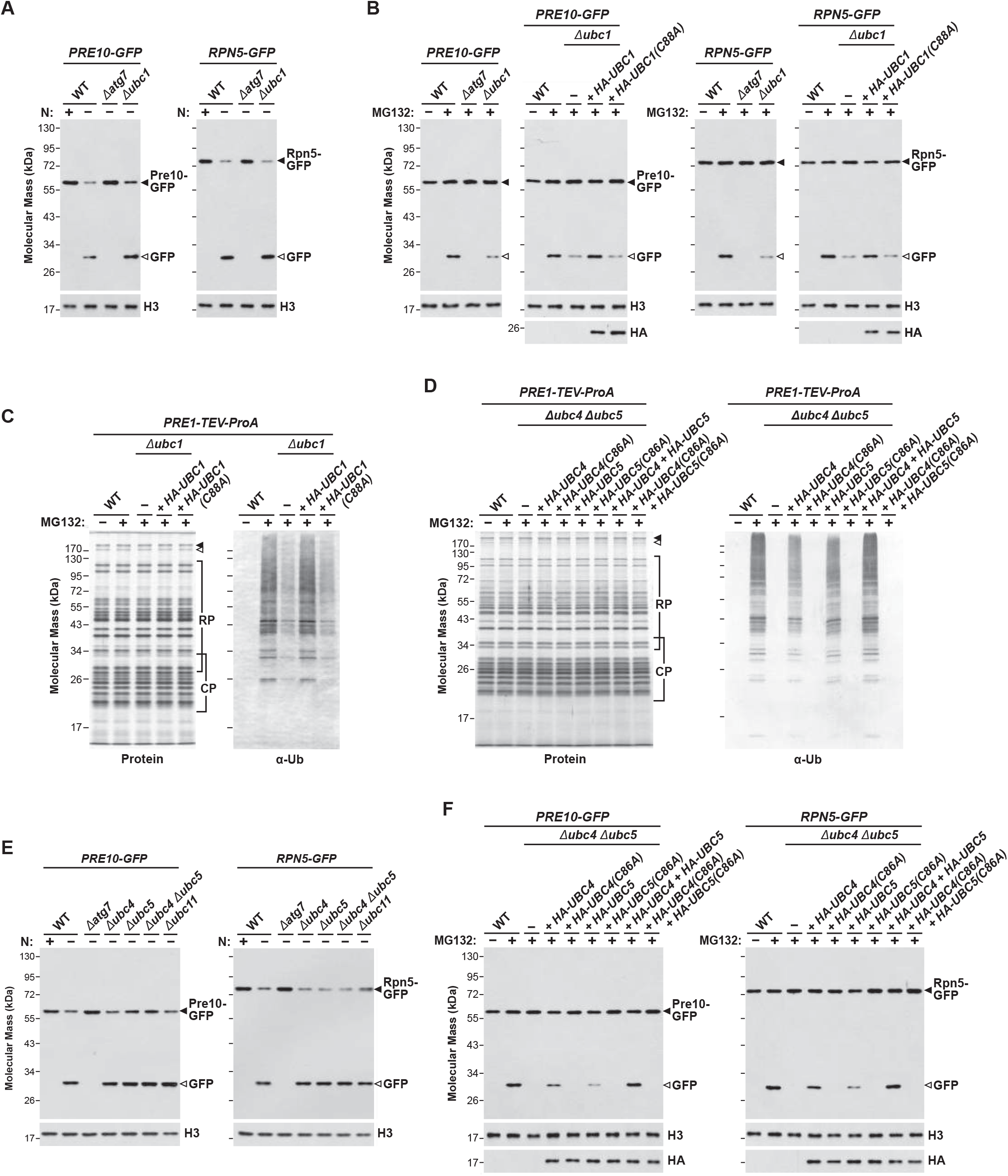
Full Description of the Impacts of Ubc1, Ubc4, and Ubc5 E2s on Ubiquitylation and Proteaphagy of Inhibited Proteasomes. Related to Figure 5. **(A)** Proteaphagy of inhibited proteasomes proceeds normally in *Δubc1* cells upon N starvation. WT and *Δubc1* cells expressing *PRE10-GFP* (left panel) or *RPN5-GFP* (right panel) were grown in +N medium at 30°C and either kept on +N medium or switched to –N medium and grown for an additional 8 hr. Cells were assayed for the release of free GFP by immunoblot analysis of cell extracts with anti-GFP antibodies. Immunodetection of histone H3 was included to confirm near equal protein loading. The *Δatg7* mutant was included as a positive control. **(B)** Proteaphagy of inhibited proteasomes requires an enzymatically active Ubc1 E2. WT and *Δubc1* cells expressing *PRE10-GFP* (left panels) or *RPN5-GFP* (right panels), either alone or complemented with HA-tagged versions of wild-type Ubc1 or the active site mutant Ubc1(C88A), were grown for 8 hr in +N medium at 30°C with or without 80 µM MG132. Cells were assayed for release of free GFP as in (A). Immunoblot analysis with anti-HA antibodies confirmed expression of the HA-tagged Ubc1 variants. **(C)** Enzymatically active Ubc1 is required for the ubiquitylation of inhibited proteasomes. Proteasomes were affinity purified from *PRE1-TEV-ProA* cells, either WT or harboring the *Δubc1* mutation alone or complemented with wild-type Ubc1 or the active site Ubc1(C88A) mutant, that were pre-treated with or without MG132 for 8 hr. The samples were subjected to SDS-PAGE and either stained for protein or immunoblotted for Ub. Migration positions of the CP and RP subunits are shown by the brackets. Closed and open arrowheads locate the Blm10 and Ecm29 accessory proteins, respectively. **(D)** Enzymatically active Ubc4 and Ubc5 are required for the ubiquitylation of inhibited proteasomes. Proteasomes were affinity purified from *PRE1-TEV-ProA* cells, either WT or harboring the *Δubc4 Δubc5* double mutations alone or complemented with wild-type Ubc4, Ubc5, and/or their corresponding active site Ubc4(C86A) or Ubc5(C86A) mutants, that were pre-treated for 8 hr with or without MG132. The samples were subjected to SDS-PAGE and either stained for protein or immunoblotted for Ub as in (C). **(E)** Proteaphagy proceeds normally during N starvation in *Δubc4* and *Δubc5* mutants. WT, *Δubc4*, *Δubc5*, and *Δubc4 ubc5* cells expressing *PRE10-GFP* (left panel) or *RPN5-GFP* (right panel) were grown on +N medium at 30°C and either kept on +N medium or switched to –N medium and grown for an additional 8 hr. Cells were assayed for the release of free GFP as in (A). The *Δatg7* mutant and the *Δubc11* mutant impacting the unrelated E2 Ubc11 were included as positive and negative controls, respectively. **(F)** Proteaphagy of inhibited proteasomes requires enzymatically active Ubc4 and Ubc5. WT and *Δubc4 Δubc5* cells expressing *PRE10-GFP* (left panels) or *RPN5-GFP* (right panels) either alone or complemented with HA-tagged versions of wild-type Ubc4 or Ubc5 or the active site mutants Ubc4(C86A) and Ubc5(C86A) were grown for 8 hr in +N medium at 30°C with or without MG132. Cells were assayed for the release of free-GFP as in (A). Immunoblotting with anti-HA antibodies confirmed expression of the HA-tagged Ubc4 and Ubc5 variants.

**Figure S7.**
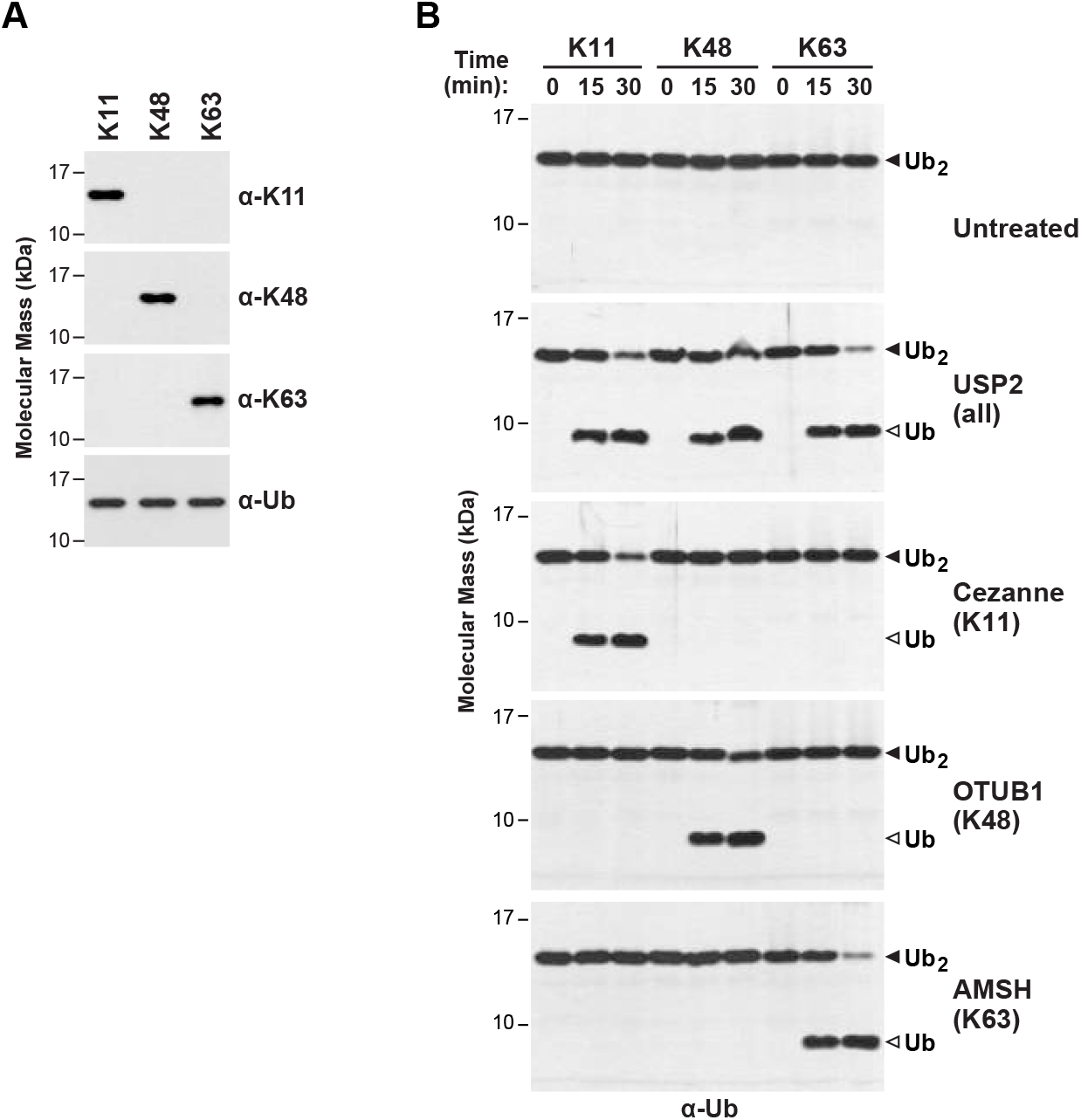
Confirmation of the Linkage-Specificity of the Ub Antibodies and DUBs Used in This Study. Related to Figure 7. **(A)** Immunoblot analysis of K11-, K48-, and K63-linked Ub-Ub dimers with anti-K11, anti-K48 and anti-K63 linkage-specific antibodies. The antibodies were used to probe equal amounts (250 ng) of Ub dimers connected by K11, K48 and K63 isopeptide linkages. Immunoblotting with general anti-Ub antibodies confirmed near equal protein loading. **(B)** The non-specific DUB USP2, and the linkage-specific DUBs Cezanne, OTUB1 and AMSH that prefer K11, K48 and K63 Ub-Ub linkages, respectively, were verified for their ability to selectively cleave K11-, K48- and K63-linked Ub dimers. Equal concentrations (10 nM) of each DUB (in 300 µl) were incubated for 0, 15 and 30 min at 30°C with 25 µg of Ub dimer internally connected via K11, K48 and K63 isopeptide bonds through the C-terminal glycine of the proximal Ub. The digestion products were subjected to SDS-PAGE and stained for total protein with silver. Closed and open arrowheads locate the Ub dimer and the released monomer, respectively.

